# ZFP541 is indispensable for pachytene progression by interacting with KCTD19 and activates meiotic gene expression in mouse spermatogenesis

**DOI:** 10.1101/2021.06.22.449524

**Authors:** Yushan Li, Ranran Meng, Shanze Li, Bowen Gu, Xiaotong Xu, Haihang Zhang, Tianyu Shao, Jiawen Wang, Yinghua Zhuang, Fengchao Wang

## Abstract

Meiosis is essential for fertility in sexually reproducing species, extensive studies tried to delineate this sophisticated process. Notwithstanding, the molecules involved in meiosis have not been fully characterized. In this study, we investigate the role of zinc finger protein 541 (ZFP541) and its interacting protein potassium channel tetramerization domain containing 19 (KCTD19) in mice. We demonstrate that they are indispensable for male fertility by regulating proper pachytene progression. ZFP541 is expressed starting from leptotene to round spermatids, and KCTD19 is initially expressed in pachytene. Depletion of *Zfp541* or *Kctd19* leads to infertility in male mice, and exhibits retarded progression from early to mid/late pachynema. In addition, *Zfp541^-/^*^-^ spermatocytes show abnormal programmed DNA double-strand breaks (DSBs) repair, impaired crossover formation/resolution, and asynapsis of the XY chromosomes. Immunoprecipitation-mass spectrometry (IP-MS) and *in vitro* Co-IP reveal that ZFP541 interacts with KCTD19, histone deacetylase 1/2 (HDAC1), HDAC2 and deoxynucleotidyltransferase terminal-interacting protein 1 (DNTTIP1). Furthermore, RNA-seq and CUT&Tag analyses demonstrate that ZFP541 binds to the promoter regions of genes involved in meiosis and post-meiosis including *Kctd19*, and activates their transcription. Taken together, our studies reveal a ZFP541-*Kctd19* transcription regulatory axis and the crucial role of ZFP541 and KCTD19 for pachytene progression and fertility in male mice.

## Introduction

Meiosis, characterized with a single round of DNA replication followed by two rounds of chromosome segregation, is a specialized cell division that generates haploid gametes for sexual reproduction and fosters genetic diversity (Gerton and Hawley, 2005; Luangpraseuth-Prosper et al., 2015). Meiosis prophase I is a relatively long period during which a series of orderly events happen. During meiotic prophase I, homologous chromosomes pair and synapsis, creating a context that promotes formation of crossover recombination events (Baker et al., 1996; Guan et al., 2020; Zickler and Kleckner, 2015). When autosomes complete synapsis at early pachynema, the heterologous XY chromosomes only pair at a short pseudoautosomal region (PAR), and the remainder of the XY chromosomes remain unsynapsed which are compartmentalized into a nuclear subdomain termed the XY body or sex body (Burgoyne et al., 2009; Li et al., 2021; McKee and Handel, 1993). In response to asynapsis, the XY chromosomes undergo chromatin modifications resulting in rapid transcriptional silencing of XY-linker genes in a process, known as meiotic sex chromosome inactivation (MSCI) which persists throughout pachytene and diplotene stages (Abe et al., 2020; Ichijima et al., 2012; Turner, 2007, 2015). A successful synapsis of autosomes and a partial synapsis of sex chromosomes are indispensable for DNA repair, recombination and subsequent desynapsis in order to ensure proper disjunction at metaphase I (Burgoyne et al., 2009; Moens, 1994). Over the past decades, many biomolecules involved in meiotic prophase I have been widely studied. Numerous knockout mouse models have been generated for studying the events during meiotic prophase I (Baudat et al., 2013; Handel and Schimenti, 2010; Jamsai and O’Bryan, 2011; Matzuk and Lamb, 2008; Roeder and Bailis, 2000). However, the machineries regulating this progression are not fully characterized (Subramanian and Hochwagen, 2014; Zickler and Kleckner, 2015).

ZFP541, also known as SHIP1, has been previously identified in the spermatocyte UniGene library, which displays testis-specific expression (Choi et al., 2008). ZFP541 contains five zinc finger motifs that bind to DNA, and an ELM2-SANT domain which recruits and activates HDAC1/2 (Choi et al., 2008; Mondal et al., 2020). Choi *et al*. (2008) found that ZFP541 forms a complex with KCTD19 and HDAC1 in adult testes, and valproic acid (VPA, a HDAC inhibitor) treatment causes hyperacetylation and reduced the expression of ZFP541/KCTD19 in round spermatids, suggesting that the ZFP541/KCTD19 complex is involved in chromatin remodeling during the post-meiotic phase (Choi et al., 2008). Oura *et al*. (2021) and Horisawa-Takada *et al*. (2021) have reported that the ZFP541/KCTD19 complex is indispensable for male meiosis and fertility (Horisawa-Takada et al., 2021; Oura et al., 2021). However, the specific role and the identity of its genome-wide target genes in the pachytene progression remain controversial.

In the present study, we generated *Zfp541* and *Kctd19* knockout mice using the CRISPR/Cas9 strategy to investigate ZFP541 and KCTD19 functions. Depletion of *Zfp541* and *Kctd19* led to male infertility and retarded progression from early to mid/late pachynema in mice. We also validated that ZFP541 can form protein complex with KCTD19, and regulate the gene expression responsible for meiotic and post-meiotic process. Collectively, our findings demonstrate that the ZFP541-*Kctd19* transcription regulatory axis and the ZFP541/KCTD19 complex are essential for pachytene progression and male fertility.

## Results

### *Zfp541* is expressed from leptotene spermatocytes to round spermatids in male mice

To determine the expression pattern of ZFP541 in spermatocytes, we explored *Zfp541* mRNA transcript and protein levels in testis from mice sampled at postnatal days (P) 1, 6, 10, 12, 14, 18, 21, 28, and 10-week-old (Adult). qRT-PCR analysis showed that *Zfp541* mRNA became detectable at P10, dramatically increased from P14 to P18, and then continued rising to P21 before plateauing at this level until at least 10-week-old (*Figure 1A*). To dissect the function of ZFP541 in meiosis, we developed a rabbit polyclonal antibody against ZFP541 protein. Western blotting revealed that obvious ZFP541 protein could be detected at P10, and reached its maximum detected level at P21 before plateauing at this level until at least 10-week-old (*Figure 1B*). Immunohistochemistry analysis demonstrated that ZFP541 is a nuclear protein that was expressed beginning in leptotene and zygotene spermatocytes, with high levels at pachytene, diplotene and round spermatids, but became downregulated at elongated spermatids, until it disappeared *(Figure 1-figure supplement 1)*. To gain a better observation of sub-cellular localization of ZFP541 in meiotic prophase I, we generated a *Zfp541-EGFP* transgenic mouse using CRISPR/Cas9 strategy *(Figure 1-figure supplement 2)*. Co-immunostaining for synaptonemal complex protein 3 (SYCP3) and GFP on chromosome spreads of spermatocytes from 10-week-old *Zfp541-EGFP* mice showed that ZFP541 was initially detected in leptotene and zygotene spermatocytes; by early pachynema, ZFP541 could be clearly detected throughout the chromatin of autosomes and XY chromosomes. In middle and late pachynema, ZFP541 labeling appeared as a bright signal on autosome chromatin, but the signal on XY chromosomes stayed relatively as a low level as early pachynema, this pattern persisted throughout diplonema (*Figure 1C and D*). These results showed that ZFP541 is expressed from leptotene spermatocytes till round spermatids in male mice, with a higher expression level in autosomes than in XY chromosomes.

**Figure 1.**
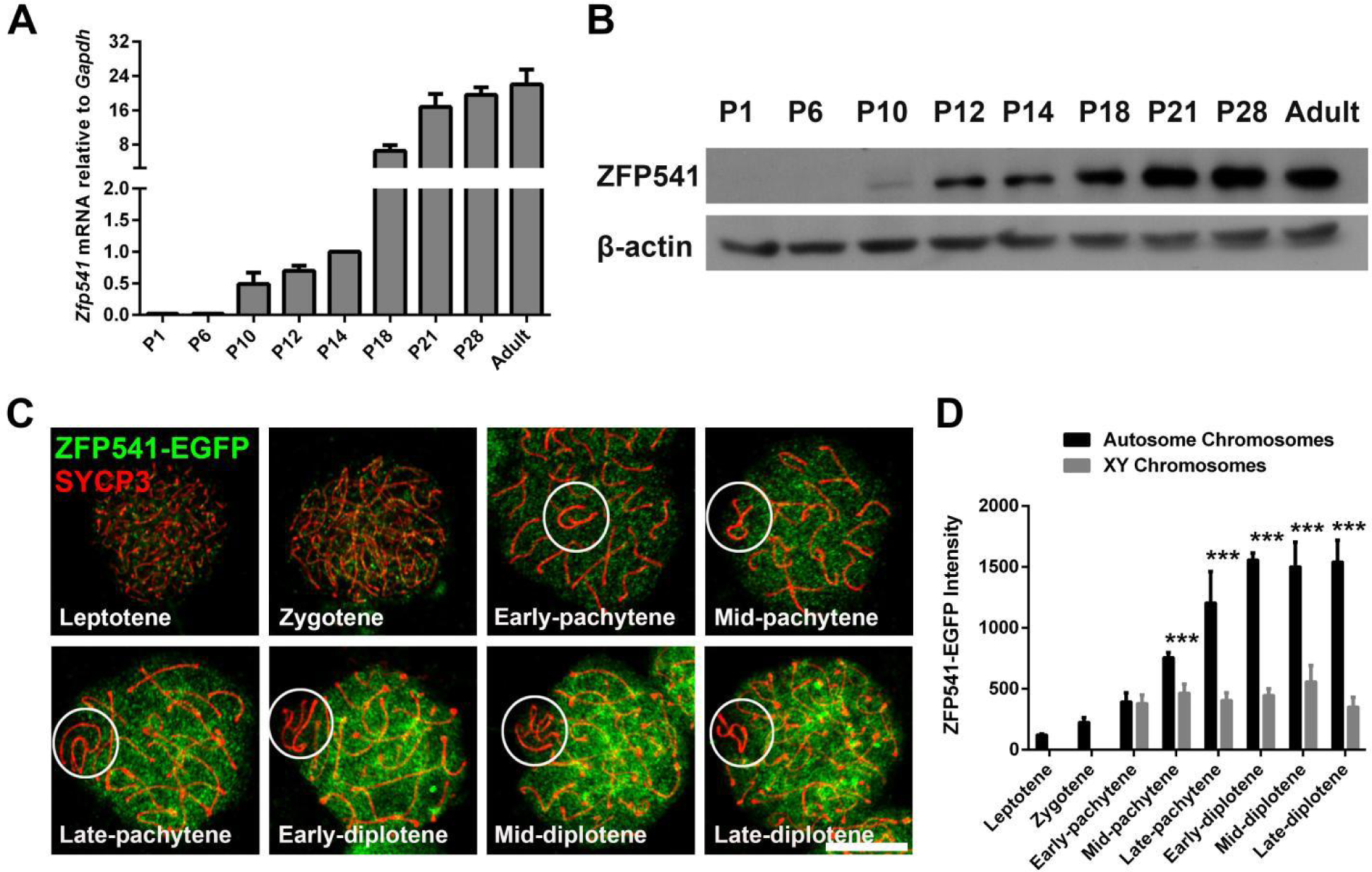
Expression Pattern of *Zfp541* in mouse testes. **(A)** qRT-PCR analysis of the relative expression of *Zfp541* mRNA obtained from mouse testes collected at different time points during postnatal development, all results were normalized to levels of *Gapdh.* (n = 3, Student’s *t*-test). **(B)** Western blotting analysis of ZFP541 in mouse testes lysates collected at different time points during postnatal development; β-actin was used as a loading control. **(C)** Immunofluorescent staining of SYCP3 on chromosome spreads of adult *Zfp541-EGFP* males. SYCP3 staining (red); ZFP541-EGFP staining (green). XY bodies are indicated by circles. Scale bars: 10 μm. **(D)** Signal intensity quantification of ZFP541 in autosomes or XY chromosomes during each stage of meiotic prophase I (n = 3, *\*\*\*P* < 0.001, Student’s *t*-test).

### *Zfp541* is essential for male fertility and pachytene progression

To elucidate the *in vivo* functions of *Zfp541*, we used the CRISPR/Cas9 strategy to generate a *Zfp541* knockout mouse model by targeted deletion of exon 3, and successfully confirmed the inactivation of this gene at the genome, mRNA transcription, as well as protein expression levels (*Figure 2-figure supplement 1 A-D*). Although developed normally, *Zfp541*^-/-^ males were sterile, in agreement with previous studies (Horisawa-Takada et al., 2021; Oura et al., 2021). The testes, the testis to body weight ratio and the seminiferous tubule diameter were all significantly smaller in the adult *Zfp541*^-/-^ males than *Zfp541*^+/-^ males (*Figure 2-figure supplement 1 E-G*). Histological examination showed that seminiferous tubules of *Zfp541^+/-^* testes contained all stages of differentiated spermatocytes as well as round and elongated spermatids, whereas *Zfp541^-/-^* testes displayed no round or elongated spermatids (*Figure 2A*). In addition, no mature spermatozoa were found in the cauda epididymis of *Zfp541^-/-^* mice (*Figure 2B*). These results suggested that ZFP541 is essential for mouse spermatogenesis and male fertility.

**Figure 2.**
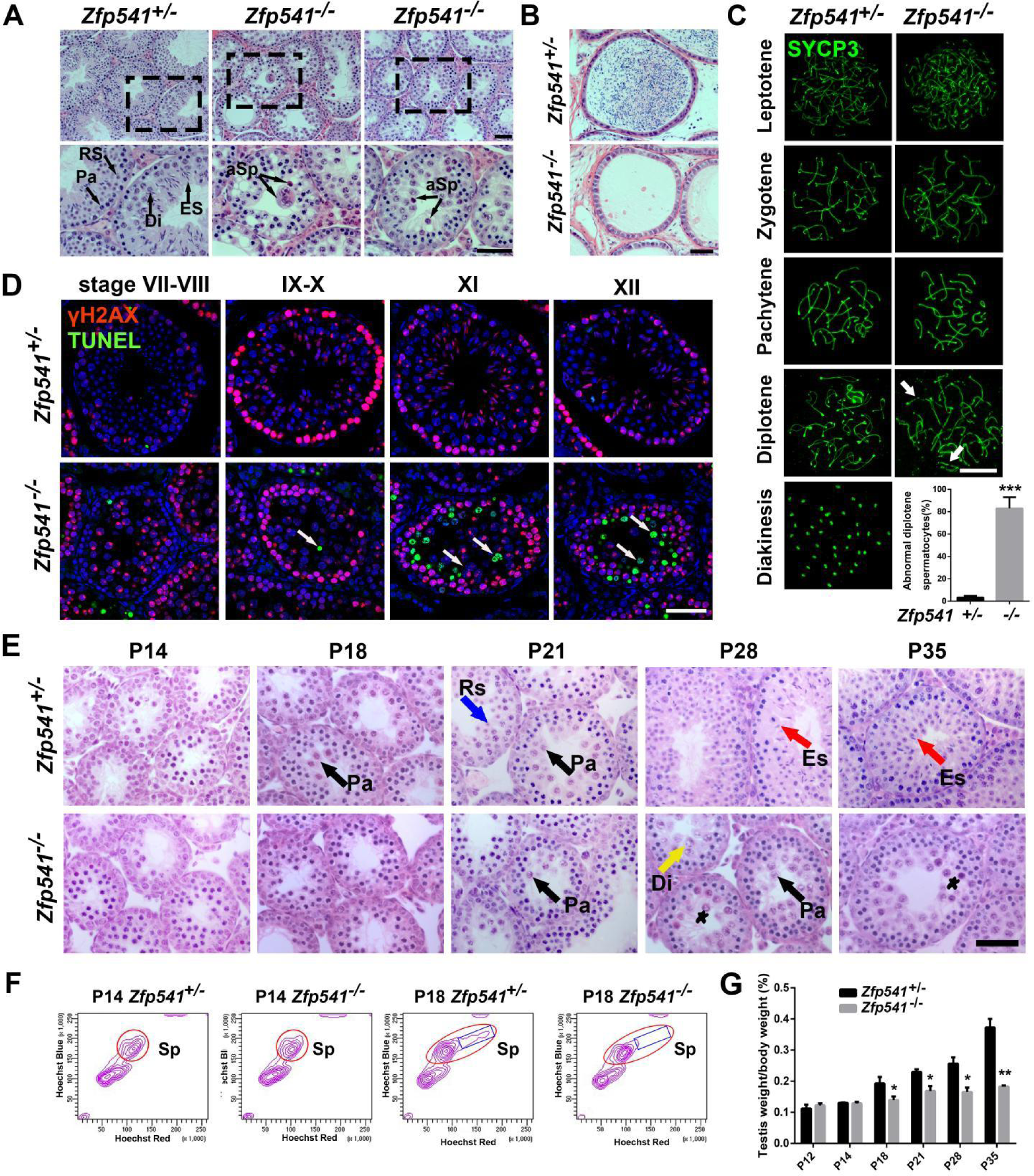
The meiotic prophase I is disrupted and arrested at the diplotene stage in the *Zfp541*^−/−^ males. **(A)** Hematoxylin and eosin staining of testes from 10-week-old *Zfp541*^+/-^ and *Zfp541*^−/−^ males. Pachytene (Pa), Diplotene (Di), Round spermatids (Rs), elongating spermatids (ES), abnormal spermatocytes (aSp). Scale bars: 50 μm. **(B)** Hematoxylin and eosin staining of caudal epididymides from 10-week-old *Zfp541*^+/-^ and *Zfp541*^−/−^ males. Scale bars: 50 μm. **(C)** Chromosome spreads of spermatocytes in testes of 10-week-old *Zfp541*^+/-^ and *Zfp541*^−/−^ males immunostained by SYCP3. Arrows indicates chromosome breakage of diplotene spermatocytes. Scale bars: 10 μm. The percentage of abnormal chromosomes in *Zfp541*^+/-^ and *Zfp541*^−/−^ diplotene spermatocytes (n= 3, *\*\*\*P* < 0.001, Student’s *t*-test). **(D)** Immunofluorescent staining of TUNEL and γH2AX in testes from 10-week-old *Zfp541*^+/-^ and *Zfp541*^−/−^ males. TUNEL staining (green); γH2AX staining (red); DAPI nuclear counterstaining of DNA (bule). Arrows indicate apoptotic spermatocytes. Scale bars: 50 μm. **(E)** Hematoxylin and eosin staining of seminiferous tubules from mouse testes collected at different postnatal stages of *Zfp541*^+/-^ and *Zfp541*^−/−^ males. Pachytene (Pa), Diplotene (Di), Round spermatids (Rs), Elongating spermatids (ES). Asterisks indicates abnormal spermatocytes. Scale bars: 50 μm. **(F)** Hoechst 33342 and PI fluorescence flow analysis of testicular cells prepared from P14 and P18 *Zfp541*^+/-^ and *Zfp541*^−/−^ males. Cells are visualized in a “Hoechst Blue”/“Hoechst Red” contour plot. Circles indicate meiotic spermatocytes; rectangle boxes indicate the shift of red Hoechst fluorescence of spermatocytes I during meiosis. **(G)** The testis to body weight ratio at different postnatal stages of *Zfp541*^+/-^ and *Zfp541*^-/-^ males (n = 4, *\*P* < 0.05; *\*\*P* < 0.01, Student’s *t*-test).

To verify which stages of meiosis were blocked by *Zfp541* deletion, we immunestained SYCP3 which can identify distinct stages of meiotic prophase I. All meiotic prophase I stages can be observed in the spermatocytes of adult *Zfp541^+/-^* testes, whereas only spermatocytes from the leptotene to diplotene were present in the *Zfp541^-/-^* testes. In addition, more abnormal chromosomes were observed in *Zfp541^-/-^* diplotene spermatocytes (*Figure 2C*). Fluorescent TUNEL and Histone H2AX at serine 139 (γH2AX, a marker for distinct spermatocytes and stages of seminiferous tubules (Blanco-Rodriguez, 2009)) staining showed that there were apoptotic cells in the stage XI-XII seminiferous tubules which contained diplotene spermatocytes in the adult *Zfp541^-/-^* testes (*Figure 2D*), suggesting that *Zfp541^-/-^* spermatocytes fail to exit diplotene and are eliminated by apoptosis.

To determine whether the first wave of spermatogenesis is affected by *Zfp541* knockout, we measured the morphological changes at different developmental stages of *Zfp541^+/-^* and *Zfp541^-/-^* testes. At P14, spermatogenesis progressed to early pachytene stage in *Zfp541^+/-^* testes (Bellve et al., 1977), histological examination and flow cytometry sorting of mouse spermatocytes by Hoechst 33342 and propidium iodide (PI) staining showed no discernible abnormalities in *Zfp541^+/-^* or *Zfp541^-/-^* testes (*Figure 2E and F*). At P18, the spermatogenesis progressed to late pachytene stage in *Zfp541^+/-^* testes but not *Zfp541^-/-^* testes (*Figure 2E*), as demonstrated by a Hoechst fluorescent shift was evident in *Zfp541^+/-^* spermatocytes but not *Zfp541^-/-^* at P18 (*Figure 2F*, rectangle boxes) (Bastos et al., 2005). At P21, round spermatids were present in *Zfp541^+/-^* mice, while late pachytene spermatocytes just appeared in *Zfp541^-/-^* mice. At P28 and P35, spermatogenesis progressed to elongated spermatids in *Zfp541^+/-^* mice, while there was a large number of pachytene spermatocytes with degenerated spermatocytes remained in *Zfp541^-/-^* mice (*Figure 2E*, asterisks). Moreover, the testis to body weight ratio was significantly decreased in the *Zfp541^-/-^* testes compared with controls from P18 to P35 (*Figure 2G*). Taken together, these results indicated that ZFP541 is required for meiotic cell cycle progression from early to mid/late pachynema.

### DSBs repair, crossover and XY synapsis are perturbed in *Zfp541^-/-^* male mice

To determine whether the developmental arrest in *Zfp541^-/-^* spermatocytes was caused by defects in homologous recombination, we monitored the distribution pattern of γH2AX (a marker of DNA damage) (Neale and Keeney, 2006). We did not find significant differences between *Zfp541^+/^*^-^ and *Zfp541^-/-^* leptotene or zygotene spermatocytes. At pachytene and diplotene *Zfp541^+/-^* spermatocytes, the γH2AX signal was only present in the XY body, by contrast, abnormal γH2AX signals were retained on autosomes of *Zfp541^-/-^* pachytene and diplotene spermatocytes (*Figure 3A*, arrows), with a proportion of approximately 30% (*Figure 3B*). The γH2AX signals of testis sections in *Zfp541^-/-^* males were similar to the above results (*Figure 3-figure supplement 1*). These results indicate that DSB repair process is compromised in the *Zfp541^-/-^* spermatocytes. DSB repair in meiosis is dependent on the recruitment of specific recombination related proteins such as DMC1 (Pittman et al., 1998). We also found more DMC1 remnant foci in *Zfp541^-/-^* pachytene spermatocytes (*Figure 3C and D*), supporting malfunction of DSB repair. During recombination, DSBs are repaired by either noncrossover or crossover pathways (Guillon et al., 2005). To determine whether crossover formation was disturbed by *Zfp541* deletion, we assessed the number of the late-recombination marker mutL homolog 1 (MLH1) foci in spermatocytes. Although crossover formation in *Zfp541^-/-^* pachytene spermatocytes was readily observed, the average number of MLH1 foci in each nucleus was significantly reduced in *Zfp541^-/-^* pachytene spermatocytes compared with controls (*Figure 3E and F*), suggesting impaired crossover formation. Moreover, MLH1 foci were present in the *Zfp541^-/-^* diplotene spermatocytes, whereas nearly all MLH1 foci had disappeared in the *Zfp541^+/-^* diplotene spermatocytes (*Figure 3F*), suggesting impairment of crossover resolution. These results collectively showed that *Zfp541* plays a vital role in DSB repair and crossover formation/resolution during meiotic prophase I in male mice.

**Figure 3.**
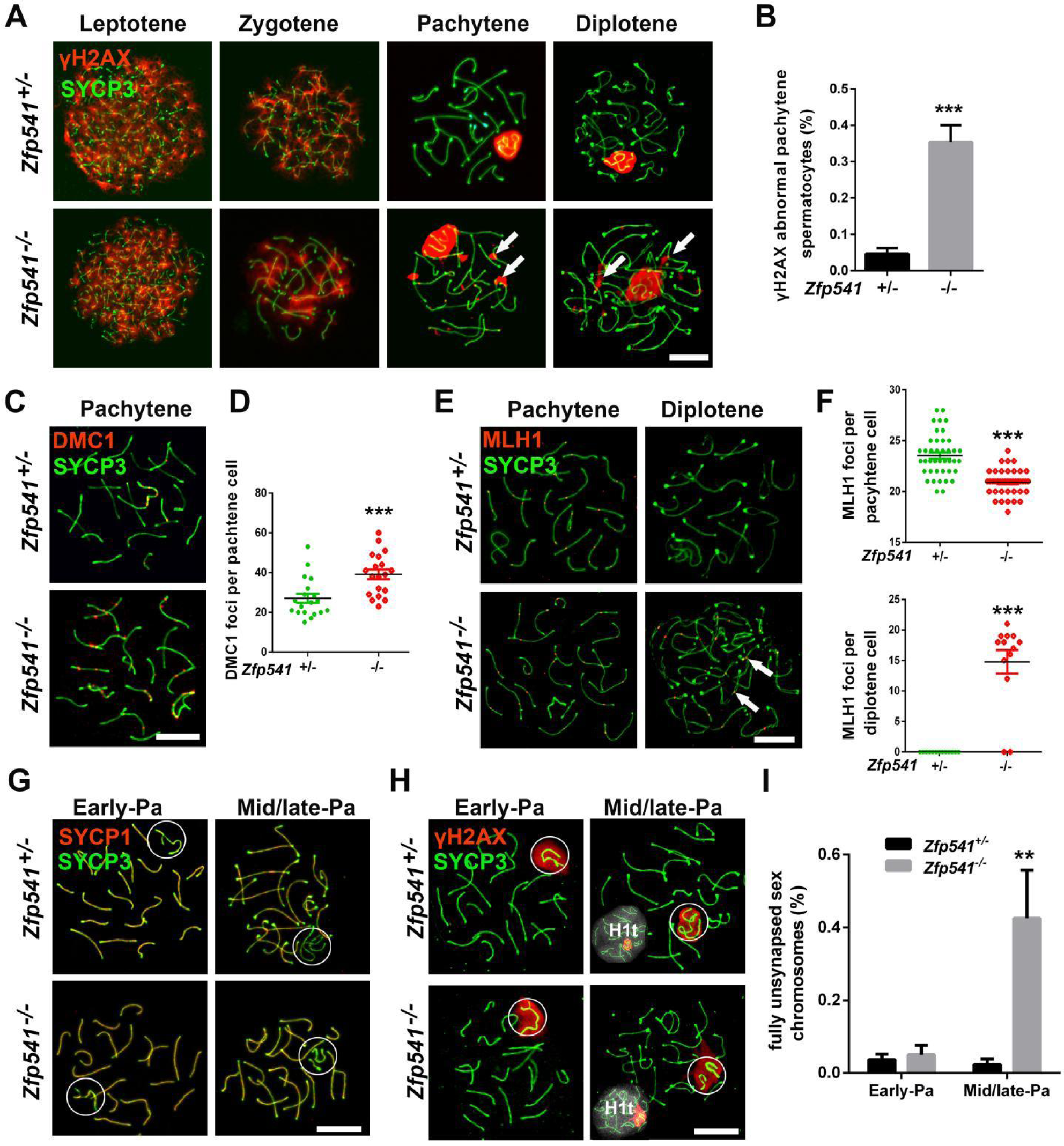
DSBs repair, crossover, and synapsis of the XY chromosomes are impaired in *Zfp541^-/-^* males. **(A)** Immunofluorescent staining of SYCP3 and γH2AX on chromosome spreads of 10-week-old *Zfp541*^+/−^ and *Zfp541*^−/−^ spermatocytes. Arrows indicate abnormal γH2AX signals retained on autosomes. SYCP3 staining (green); γH2AX staining (red). Scale bars: 10 μm. **(B)** The percentage of pachytene spermatocytes retained γH2AX in the autosomal region in the 10-week-old *Zfp541*^+/-^ and *Zfp541*^−/−^ males (n = 4, *\*\*\*P* < 0.001, Student’s *t*-test). **(C)** Immunofluorescent staining of SYCP3 and DMC1 on chromosome spreads of *Zfp541*^+/-^ and *Zfp541*^−/−^ pachytene spermatocytes. SYCP3 staining (green); DMC1 staining (red). Scale bars: 5 μm. **(D)** Quantification of DMC1 foci per cell in *Zfp541*^+/−^ and *Zfp541*^−/−^ pachytene spermatocytes (n = 4, *\*\*\*P* < 0.001, Student’s *t*-test). **(E)** Immunofluorescent staining of SYCP3 and MLH1 on chromosome spreads of *Zfp541*^+/-^ and *Zfp541*^−/−^ pachytene and diplotene spermatocytes. Arrows show MLH1 foci on the *Zfp541*^−/−^ diplotene spermatocytes. SYCP3 (green); MLH1 staining (red). Scale bars: 10 μm. **(F)** Quantification of MLH1 foci per cell in *Zfp541*^+/−^ and *Zfp541*^−/−^ pachytene and diplotene spermatocytes (n= 4, *\*\*P* < 0.01, Student’s *t*-test). Scale bars: 10 μm. **(G)** Immunofluorescent staining of SYCP3 and SYCP1 on chromosome spreads of *Zfp541*^+/−^ and *Zfp541*^−/−^ early and mid/late pachytene (Pa) spermatocytes. SYCP3 staining (green); SYCP1 staining (red). Scale bars: 10 μm. **(H)** Immunofluorescent staining of SYCP3, γH2AX, and H1t on chromosome spreads of *Zfp541*^+/-^ and *Zfp541*^−/−^ early and mid/late pachytene (Pa) spermatocytes. SYCP3 staining (green); γH2AX staining (red); H1t staining (white). Scale bars: 10 μm. **(I)** Quantification of the XY chromosomes with synapsis defects at the PAR in *Zfp541*^+/−^ and *Zfp541*^−/−^ pachytene spermatocytes (n = 4, *\*\*P* < 0.01, Student’s *t*-test).

To investigate whether failure to resolve DSBs and to form crossovers in *Zfp541^-/-^* spermatocytes was caused by defects in synapsis, we next analyzed the dynamic pattern of chromosome pairing and synapsis by monitoring SYCP3 and synaptonemal complex protein 1 (SYCP1) location. Chromosomes of both *Zfp541^+/-^* and *Zfp541^-/-^* early pachytene spermatocytes were synapsed, while *Zfp541^-/-^* mid/late pachytene spermatocytes exhibited asynapse of XY chromosomes (*Figure 3G*). To quantify the timing of this defect, we divided pachytene-stage cells into early and mid/late pachynema by immunostaining for the testis-specific histone H1 variant (H1t) (Drabent et al., 1996). In early spermatocytes (H1t negative), the proportion of the XY chromosome synapsis was indistinguishable in *Zfp541^+/-^* and *Zfp541^-/-^* spermatocytes (*Figure 3H*). However, in spermatocytes that had progressed beyond mid/late pachynema, the proportion of fully asynapsis of XY chromosomes (H1t positive) was significantly increased in the *Zfp541^-/-^* spermatocytes (*Figure 3I*). These data showed that synapsis of terminal XY chromosomes is normally reinforced by nascent crossing-over between the PAR in the early pachynema, and that the recombination defect of *Zfp541^-/-^* mutants can lead to premature asynapsis of XY chromosomes during mid/late pachynema.

Given the asynapsis of XY chromosomes in *Zfp541^-/^*^-^ testes, we wonder whether characteristic histone modifications on XY chromosomes might be affected in the absence of Z*fp541* (Abe et al., 2020). H3 trimethylated at lysine 9 (H3K9me3), the histone mark associated with chromatin condensation and transcriptional repression (Page et al., 2012), was localized in the XY chromosomes of early pachynema, and disappeared in late pachynema, but in Zf*p541^-/^*^-^ spermatocytes, H3K9me3 was not removed properly from the Y chromosome at late pachynema (*Figure 3-figure supplement 2A*). Another maker for transcriptionally active chromatin (Page et al., 2012), histone H3 acetylated at lysine 9 (H3K9ac) is gradually established from early pachynema to late pachynema, but we did not observe the H3K9ac signal in the Y chromosome of *Zfp541^-/^*^-^ late pachynema spermatocytes (*Figure 3-figure supplement 2B*), suggesting that the abnormal expression of H3K9me3 and H3K9ac in the Y chromosome may lead to asynapsis of XY chromosomes from middle pachytene to early diplotene spermatocytes in the *Zfp541^-/-^* males. Collectively, our data indicated that ZFP541 is required for the synapsis of XY chromosomes from middle pachynema to diplonema.

### *Kctd19* is required for male fertility by regulating progression from early to mid/late pachynema

KCTD19 associates with ZFP541 and HDAC1 and is required for meiotic exit in male mice (Horisawa-Takada et al., 2021; Oura et al., 2021). We wonder if KCTD19 is also vital for spermatogenesis, so we explored the expression pattern of KCTD19. qRT-PCR assays showed that *Kctd19* mRNA was at very low levels at P10 and P12, dramatically increased from P14 to P18, and then continued rising to P21 before plateauing at this level until at least 10-week-old. Western blotting and immunohistochemistry (a rabbit polyclonal antibody against KCTD19 of our lab) showed that KCTD19 protein was specifically localized in the cell nuclei from pachytene spermatocytes to round spermatids (*Figure 4-figure supplement 1*). To gain a better observation of sub-cellular localization of KCTD19 in meiotic prophase I, we generated *Kctd19-EGFP* mice (*Figure 4-figure supplement 2*). Immunostaining for SYCP3 and GFP on chromosome spreads of *Kctd19-EGFP* spermatocytes showed that KCTD19 was not observed in the nuclei of leptotene or zygotene spermatocytes. It was first detected in early pachytene spermatocytes with a widespread localization in the whole nuclei, and this pattern persisted to the diplotene stage. Also, well-defined and condensed KCTD19 signals were shown in the XY body area from early pachytene to early diplotene stage (*Figure 4A*). Collectively, these data showed that KCTD19 is expressed starting from pachytene spermatocytes.

**Figure 4.**
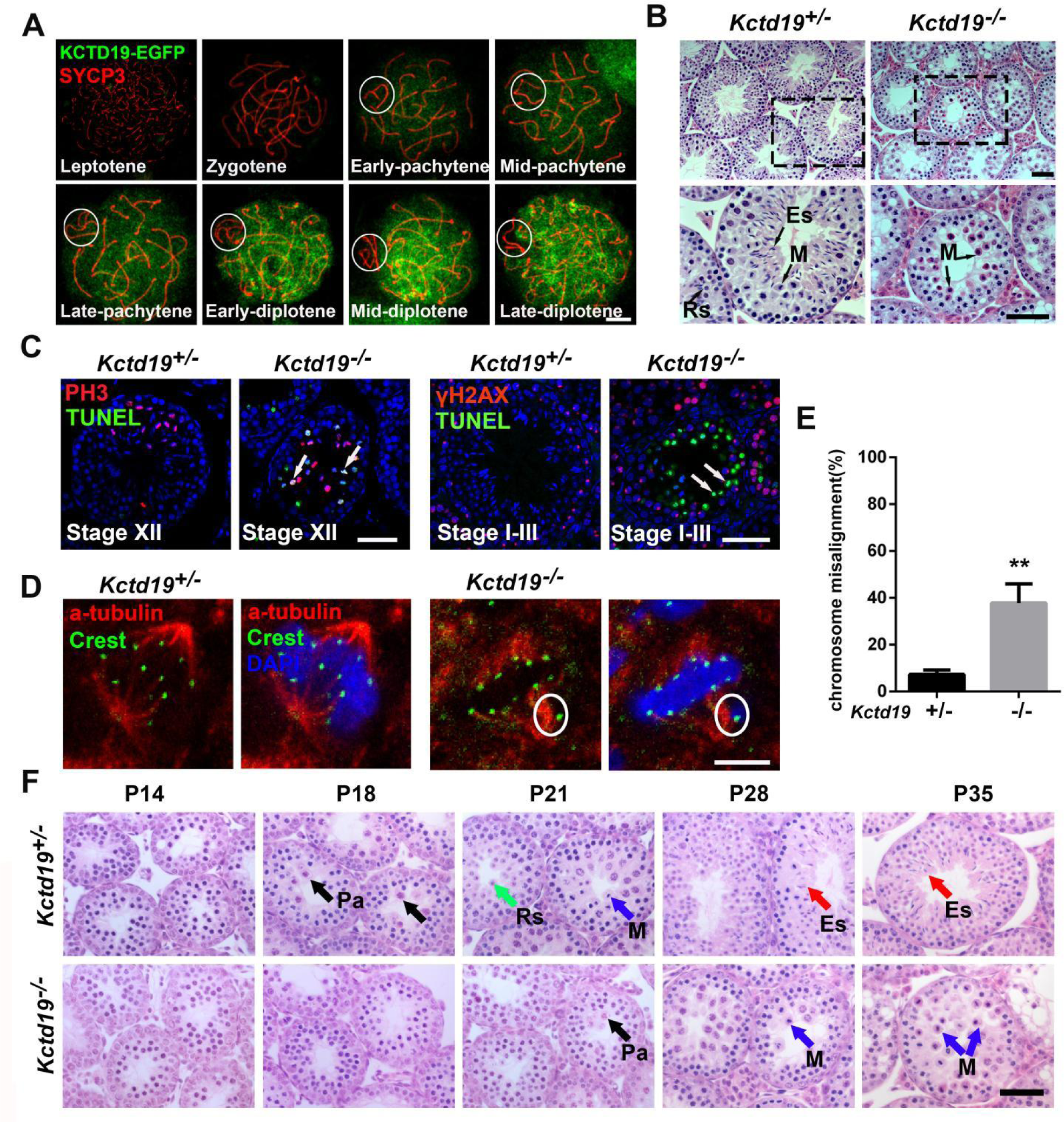
Spermatogenesis arrests at meiotic metaphase in *Kctd19*^−/−^ males. **(A)** Immunofluorescent staining of SYCP3 and KCTD19 co-expression on chromosome spreads of 10-week-old *Kctd19-EGFP* males. SYCP3 (red); KCTD19-GFP (green). White circles show XY bodies. Scale bars: 10 μm. **(B)** Hematoxylin and eosin staining of mouse testes from 10-week-old *Kctd19*^+/-^ and *Kctd19*^−/−^ males. Round spermatids (Rs), elongating spermatids (ES), Metaphase (M). Scale bars: 50 μm. **(C)** Immunofluorescent staining of TUNEL, PH3 or γH2AX in *Kctd19*^+/-^ and *Kctd19*^−/−^ testes. Arrows indicate apoptotic cells. TUNEL staining (green); PH3 and γH2AX staining (red). Scale bars: 50 μm. **(D)** Immunofluorescent staining of CREST and α-TUBULIN between 10-week-old *Kctd19*^+/-^ and *Kctd19*^−/−^ testes. CREST staining (green); α-TUBULIN staining (red); DAPI nuclear counterstaining of DNA (bule). White circles indicate misaligned chromosomes. Scale bars: 5 μm. **(E)** The percentage of metaphase I spermatocytes with misaligned chromosomes between 10-week-old *Kctd19*^+/-^ and *Kctd19*^−/−^ testes (n = 4, *\*\*P* < 0.01, Student’s *t*-test). **(F)** Hematoxylin and eosin staining of mouse testes collected at different postnatal stages of *Kctd19*^+/-^ and *Kctd19*^−/−^ males. Pachytene (Pa), Metaphase (M), Round spermatids (Rs), Elongating spermatids (ES). Scale bars: 50 μm.

To characterize the *in vivo* functions of *Kctd19*, we then generated *Kctd19^-/-^* mice with targeted deletion at exon 2, and confirmed the inactivation of this gene (*Figure 4-figure supplement 3A-D*). Similar to *Zfp541*^-/-^ mice, *Kctd19*^-/-^ males were completely infertile. The testes, the testis to body weight ratio, and the seminiferous tubule diameter were all significantly smaller in the 10-week-old *Kctd19*^-/-^ males than controls (*Figure 4-figure supplement 3E-G*). Histological examination showed that *Kctd19^-/-^* males displayed no round or elongated spermatids in the seminiferous tubules, and arrested at meiotic metaphase (*Figure 4B*). In addition, no mature spermatozoa were found in the cauda epididymis of the *Kctd19*^-/-^ mice (*Figure 4-figure supplement 3H*). Fluorescent TUNEL, phosphorylated histone H3 (PH3, a characteristic marker of the transition into metaphase) or γH2AX staining revealed that metaphase spermatocytes underwent apoptosis in the seminiferous stage XII, and the number of apoptotic spermatocytes accumulated in the seminiferous stage I-III tubules (*Figure 4C*). We immune-stained SYCP3, γH2AX, SYCP1, and MLH1 in meiosis, but found no obvious alteration in DSB repair, synapse or crossover in *Kctd19*^−/−^ spermatocytes (*Figure 4-figure supplement 4*), indicating that homologous recombination process is not affected by *Kctd19*. Nevertheless, immunostaining for CREST (centromere marker) and α-TUBULIN (microtubule marker) in frozen sections revealed chromosome misalignment on the metaphase plate in the 10-week-old *Kctd19*^−/−^ testes (*Figure 4D and E*). Altogether, these results demonstrated that KCTD19 is required for the progression of meiotic prophase.

To determine whether the first wave of spermatogenesis is also affected by *Kctd19* knockout, we compared the morphological changes at different postnatal stages of *Kctd19^+/-^* and *Kctd19^-/-^* testes. At P14, histological analysis showed no discernible abnormality in *Kctd19^-/-^* testes as compared to controls (*Figure 4F*). At P18, the spermatogenesis progressed to late pachytene stage in *Kctd19^+/-^* testes but not *Kctd19^-/-^* testes. At P21, metaphase spermatocytes and round spermatids were present in the *Kctd19^+/-^* testes, while late pachytene spermatocytes appeared and no metaphase spermatocytes were found in *Kctd19^-/-^* testes. At P28 and P35, the spermatogenesis was arrested at the metaphase of the first meiotic division in *Kctd19^-/-^* mice (*Figure 4F*). These observations suggest that the meiotic cell cycle progression is retarded from early to mid/late pachytene stages from P14 to P18, and arrested at meiotic metaphase stage *in Kctd19^-/-^* male mice. Moreover, the two histone modification markers, H3K9me3 and H3K9ac exhibited similar pattern in *Kctd19*^-/-^ spermatocytes as those in *Zfp541^-/-^* spermatocytes (*Figure 4-figure supplement 5*). Collectively, these results showed that ZFP541 and KCTD19 are involved in regulating the developmental procession of meiotic prophase I, and the location of H3K9me3 and H3K9ac in the Y chromosome from middle pachytene to early diplotene spermatocytes.

### ZFP541 interacts with KCTD19, HDAC1, HDAC2 and DNTTIP1

In order to elucidate the function of ZFP541 and KCTD19, the interacting factors of ZFP541 and KCTD19 at P18 testes were screened by IP-MS analyses. We identified the association of ZFP541 with KCTD19, HDAC1, HDAC2 and DNTTIP1 (*Supplement file 1 and 2*), and further confirmed by immunoprecipitation of either ZFP541 or KCTD19 in the P18 *wild type* and knockout mouse testes (*Figure 5A and B*). To dissect ZFP541/KCTD19 complex protein-protein interactions, we co-transfected FLAG-tagged *Zfp541* with MYC-tagged *Kctd19, Hdac1*, *Hdac2,* and *Dnttip1* pairwise in HEK 293T cell line. The Co-IP results showed that ZFP541 interacted with KCTD19, HDAC1 DNTTIP1, and HDAC2 directly (*Figure 5C*). In addition, co-transfection with GFP-tagged *Zfp541* and MYC-tagged *Kctd19, Hdac1*, *Hdac2* and *Dnttip1* constructs to HEK 293T showed that ZFP541 co-localized with KCTD19, HDAC1, HDAC2, and DNTTIP1 in the nucleus (*Figure 5D*). These results demonstrated that ZFP541 forms a complex with KCTD19, HDAC1, HDAC2 and DNTTIP1.

**Figure 5.**
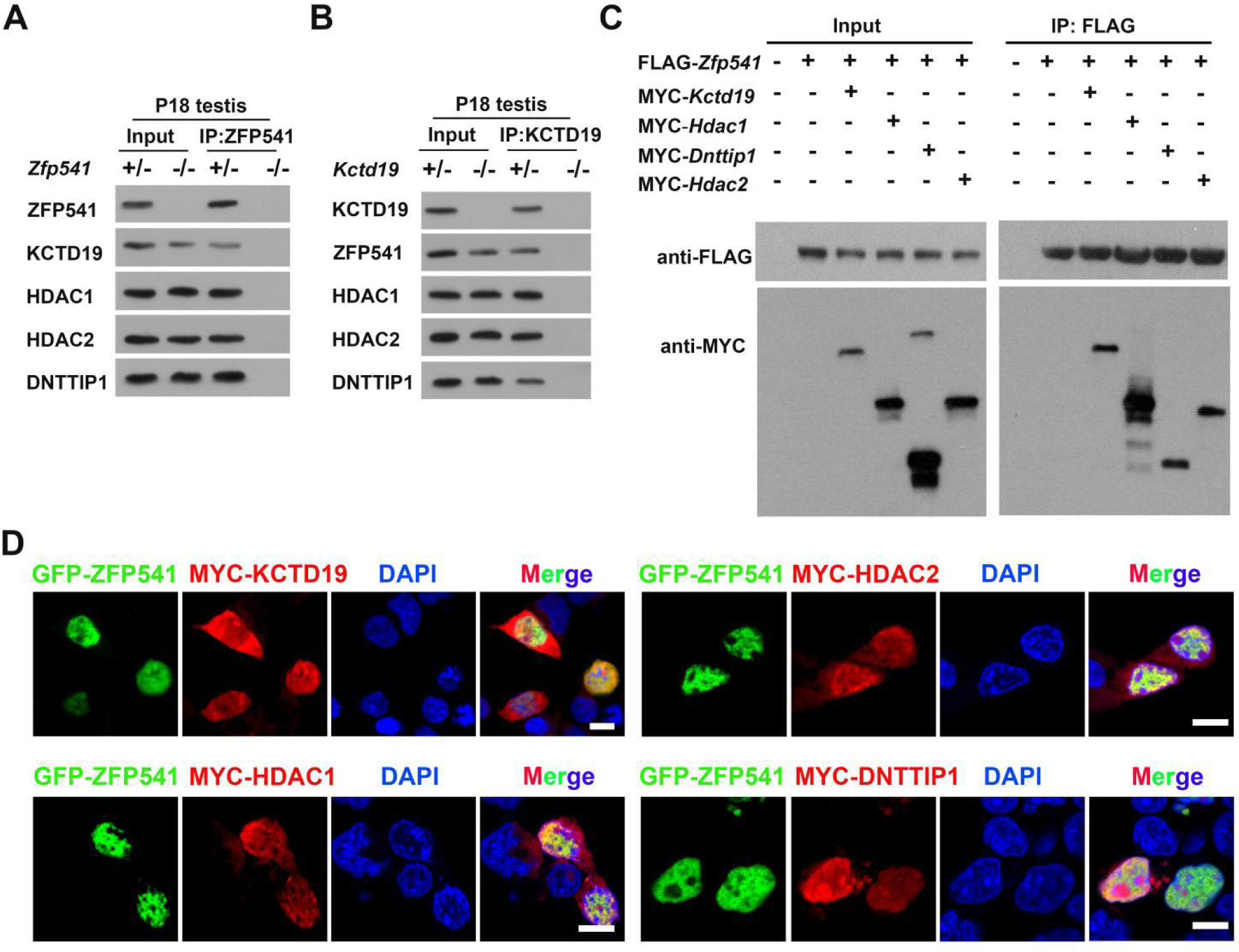
ZFP541 interacts with KCTD19, HDAC1, HDAC2 and DNTTIP1, and forms a complex. **(A)** Western blotting showing the immunoprecipitates of ZFP541 from P18 *Zfp541^+/-^* and *Zfp541^-/-^* testicular protein extracts. **(B)** Western blotting showing the immunoprecipitates of KCTD19 from P18 *Kctd19^+/-^* and *Kctd19^-/-^* testicular protein extracts. **(C)** FLAG-tagged and/or MYC-tagged expression constructs were transfected into HEK 293T cells, followed by Co-IP with anti-FLAG antibody and western blotting with anti-FLAG and anti-MYC antibodies as indicated. Input was 10% of the lysate used for IP. **(D)** GFP-ZFP541 and MYC-tagged expression constructs were transfected into HEK 293T cells, followed by immunofluorescence staining with anti-GFP and anti-FLAG antibodies. Scale bars: 10 μm.

### ZFP541 regulates meiotic and post-meiotic genes expression

Due to the DNA binding affinity and interacting ability with HDACs, we wonder if ZFP541 can regulate gene expression during meiosis. We performed RNA-seq of *Zfp541*^+/-^ and *Zfp541*^-/-^ testes at P14, because *Zfp541* already started to express at this time and there were no obvious developmental defects between *Zfp541*^+/−^ and *Zfp541*^−/−^ testes. We identified 2 upregulated and 234 downregulated genes in P14 *Zfp541*^-/-^ versus *Zfp541*^+/-^ testes (fold change > 1.5, false discovery rate (FDR) < 0.05) (*Figure 6A and Figure 6-figure supplement 1*). Gene ontology (GO) analysis of the downregulated genes in *Zfp541*^-/-^ testes are mostly involved in cilium movement, spermatogenesis, fertilization, and meiotic cell cycle (*Figure 6B*). These downregulated genes of meiosis (*Kctd19*, *Hspa2, Fbxo43, Morc2b, 1700102P08Rik* (*Maps*)*, Psma8*) and post-meiosis (*Piwil1, Crisp2, Prok2, Hsf5, Adam2, Clgn, Ybx2, Spink2, Stk33, Rfx2*) were validated by qRT-PCR. At P12, expression of each of these genes had no obvious changes between *Zfp541*^+/-^ and *Zfp541*^-/-^ testes, while expression of these genes was dramatically reduced in *Zfp541*^-/-^ than *Zfp541*^+/-^ testes at P14 (*Figure 6C*). What’s more, the expression of these genes was also decreased in *Kctd19^-/-^* testes (*Figure 6-figure supplement 2*). Intriguingly, we found that the expression of both *Kctd19* mRNA and protein was significantly downregulated in *Zfp541^-/-^* testes (*Figure 6C and D*), while the expression of both *Zfp541* mRNA and protein was not affected by *Kctd19* depletion (*Figure 6E and Figure 6-figure supplement 2*). Collectively, these results demonstrated that the ZFP541/KCTD19 complex is required for the expression of spermatocyte-specific genes involved in meiotic and post-meiotic developmental processes. In addition, *Kctd19* expression is also regulated by ZFP541.

**Figure 6.**
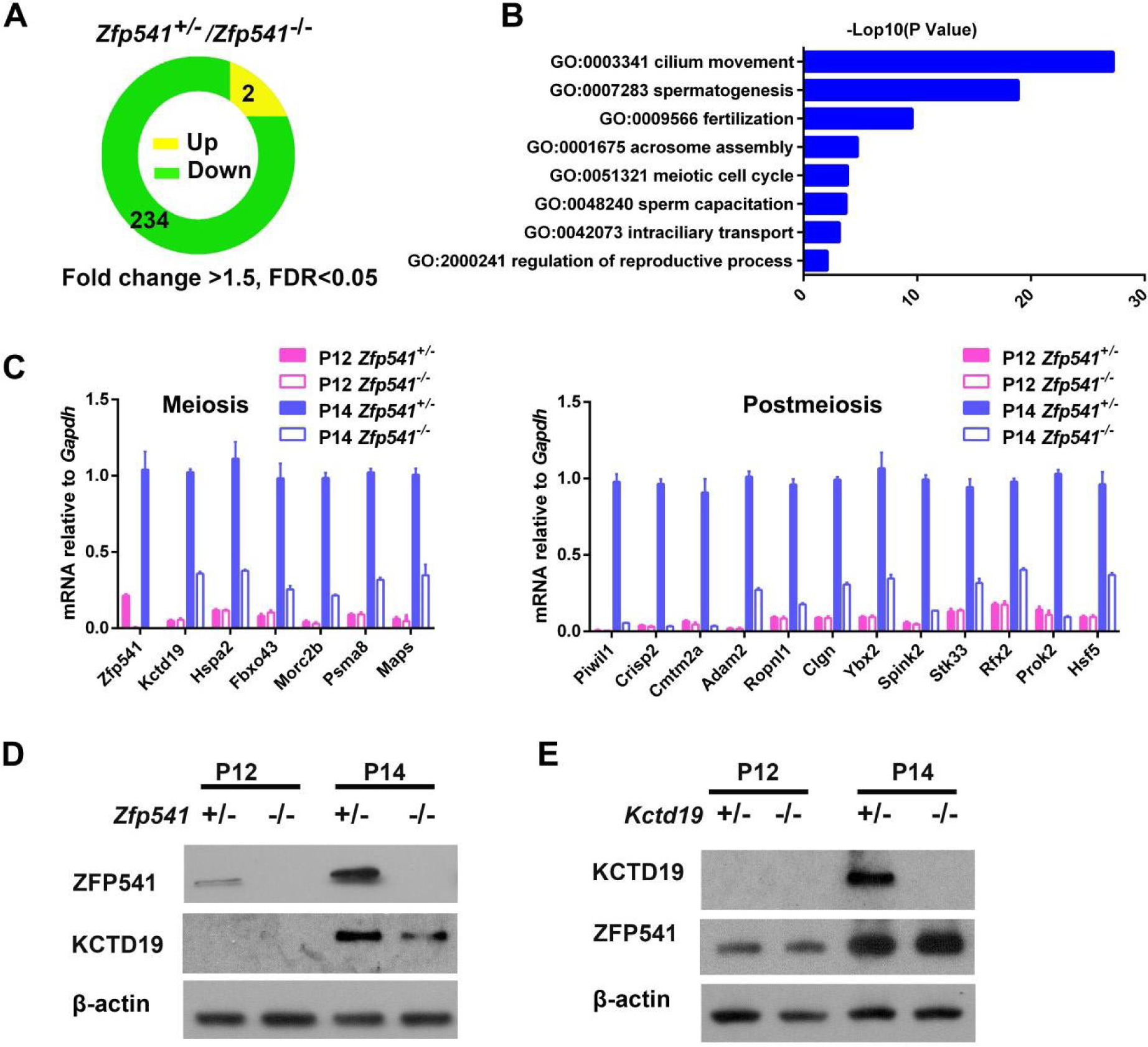
A substantial of meiosis and post-meiosis related genes are downregulated in *Zfp541*^−/−^ testes. **(A)** Genes exhibiting significant up- or down-regulation in *Zfp541*^+/−^ and *Zfp541*^−/−^ testes at P14. **(B)** GO analysis of downregulated genes in *Zfp541*^−/−^ in testis identified by RNA-seq. **(C)** Validation of RNA-seq results on expression levels of representative genes in P12 and P14 *Zfp541*^+/−^ and *Zfp541*^−/−^ testes (n = 3, Student’s *t*-test). **(D)** Western blotting analysis of ZFP541 and KCTD19 protein in P12 and P14 *Zfp541*^+/-^ and *Zfp541*^-/-^ testes. β-actin was used as a loading control. **(E)** Western blotting analysis of KCTD19 and ZFP541 protein in P12 and P14 *Kctd19*^+/-^ and *Kctd19^-/-^* testes. β-actin was used as a loading control.

### ZFP541 binds to a set of meiotic and post-meiotic genes, and activates their transcription

To determine whether ZFP541 specifically associates with its target genes and regulate their transcription, we took advantage of CUT&Tag technology to determine the enriched genome loci of ZFP541 (Kaya-Okur et al., 2019). By using P18 *wild type* and *Zfp541-EGFP* testes, when ZFP541 was abundantly expressed, we found that about 87% of the ZFP541 peaks located within the promoter regions (*Figure 7A*), and the majority of ZFP541 binding sites were close to the center of transcription start sites (TSS) (*Figure 7B*). *De novo* motif analysis of ZFP541 peaks using Multiple EM for Motif Elicitation (MEME) identified two GC rich DNA sequences as the most significantly enriched ZFP541-binding motif (*Figure 7C*). Among the ZFP541-binding targets, 127 genes were identified within the downregulated genes in *Zfp541*^−/−^ testes (*Figure 7D*). GO analysis of these genes showed that they are mostly involved in spermatogenesis and cillium movement (*Figure 7E*). Notably, ZFP541 was enriched in the promoter regions of genes involved in meiotic (*Kctd19, Hspa2, Morc2b, Maps*) and postmeiotic (*Clgn, Ybx2, Hsf5*) process (*Figure 7F*). Interestingly, strong CUT&Tag peaks were observed at the *Zfp541* promoter, indicating that ZFP541 auto-regulates its expression by targeting its own promoter (*Figure 7G*). Auto-regulation is one of the most efficient regulatory loops to maintain gene expression levels, and has also been reported for the transcription factor CREM and SOX30 that occupy their own promoter (Martianov et al., 2010; Bai et al., 2018). In contrast, ZFP541 did not bind to promoters or gene bodies of spermatogonial self-renewal *Plzf* (promyelocytic leukemia zinc-finger) or the housekeeping gene *Gapdh* (Glyceraldehyde 3-phosphate dehydrogenase) (*Figure 7H*). Collectively, these data demonstrated that ZFP541 can promote a set of meiotic and post-meiotic genes by directly binding to their promoter regions.

**Figure 7.**
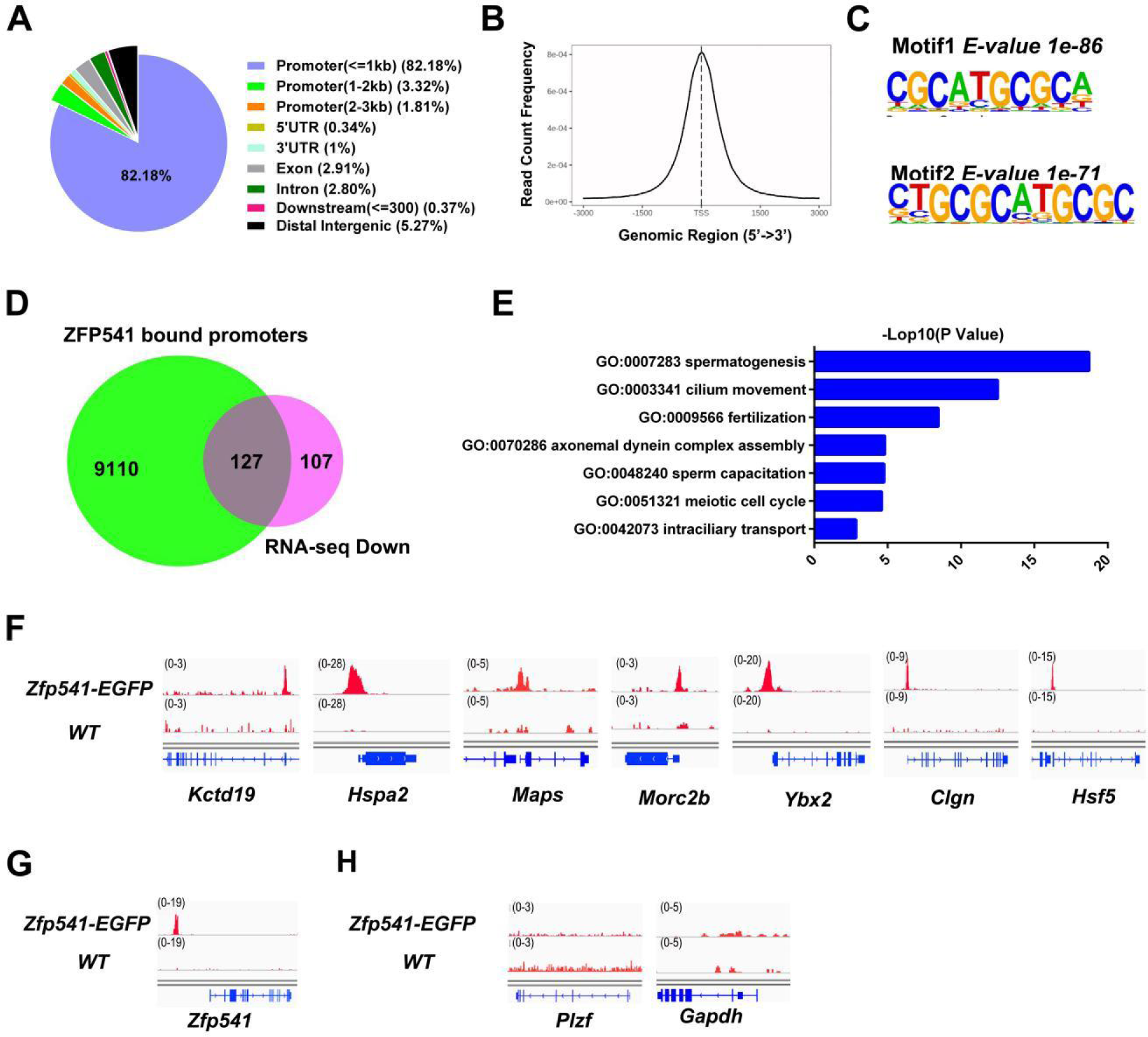
ZFP541 specifically bind and regulate expression of meiotic and post-meiotic genes. **(A)** Genomic annotation of ZFP541 binding loci. CUT&Tag experiments with anti-GFP antibody were performed using P18 *wild type* and *Zfp541-EGFP* mouse testes. **(B)** Distribution of ZFP541 CUT&Tag reads on gene bodies is plotted. ZFP541 was enriched at the regions from -1 kb to +1 kb relative to the transcription start sites (TSS). The Y axis represents the frequency of ZFP541 CUT&Tag counts at specific site, normalized by total reads counts. **(C)** *De novo* motif analysis of ZFP541 binding sites using MEME. The most enriched de novo motifs are shown. **(D)** Venn diagram representing the overlap of ZFP541-bound genes (9237 nearest genes) and downregulated (234 genes) in *Zfp541*^−/−^ mice. **(E)** GO analysis of the ZFP541-bound genes that were downregulated in *Zfp541*^−/−^ mice (127 genes). **(F)** Genome browser view of ZFP541 CUT&Tag reads on representative gene loci in testes from P18 *wild type* (*WT*) and *Zfp541-EGFP* mice. **(G)** Genome browser view of ZFP541 CUT&Tag reads on the promoter of *Zfp541*. (H) Genome browser view of ZFP541 CUT&Tag reads on the promoter of *Plzf* and the housekeeping gene *Gapdh*.

## Discussion

In this study, we demonstrate that ZFP541 forms a complex with KCTD19, HDAC1, HDAC2, and DNTTIP1 in testes, and is required for pachytene progression in meiotic prophase I. Knockout of *Zfp541* and *Kctd19* cause the retarded progression from early to mid/late pachynema, leading to apoptosis and male infertility. Our detailed phenotypic analyses showed that *Zfp541^-/-^* spermatocytes are arrested at early diplotene stage with defective DSB repair, impaired crossover, asynapsis of XY chromosomes and chromosome abnormalities of diplotene spermatocytes. It was recently reported that mouse ZFP541 and KCTD19 are co-expressed in spermatocytes from pachytene onward (Oura et al., 2021), while our results show that the expression of ZFP541 in spermatocytes starts from leptotene, a time point slightly earlier than KCTD19. The discrepancy in ZFP541 expression pattern can be attributed to higher efficacy of our antibody and the *Zfp541-EGFP* transgenic mice.

Efficient MSCI is indispensable for prophase I progression, perturbations in MSCI lead to mis-expression of sex genes and mid-pachytene spermatocyte apoptosis (Royo et al., 2010; Vernet et al., 2016). H3K9me3 is observed on the XY body at early pachytene, which is involved in MSCI initiation (Hirota et al., 2018). We found that depletion of *Zfp541* did not alter synapsis of the XY chromosomes, neither the location of H3K9me3 and H3K9ac in early pachynema. Whereas the proportion of fully asynapsis of XY chromosomes in the *Zfp541^-/-^* spermatocytes was significantly increased in the mid/late pachynema, and the locations of H3K9me3 and H3K9ac were disturbed in the Y chromosome from *Zfp541^-/-^* middle pachynema to diplotena, suggesting that *Zfp541* is required for the synapsis of XY chromosomes from middle pachynema to diplonema. To further investigate if ZFP541 affect the transcription of XY chromosome, we compared the distribution of phosphorylated RNA polymerase II (a marker of active transcription) in *Zfp541^+/-^* and *Zfp541^-/-^* pachytene and diplotene spermatocytes (*Figure 3-figure supplement 3*). Nonetheless, there is no obvious distinction in the *Zfp541^-/-^* spermatocytes. These data suggested that once MSCI was established in early pachynema, XY silencing is remarkably stable, regardless of changes in the XY synapsis and histone modifications from middle pachynema to diplonema.

Our transcriptome data show that a substantial of genes involved in spermatogenesis are regulated by ZFP541. Some of these genes are also responsible for male meiosis, such as *Kctd19*, *Morc2b* (Shi et al., 2018), *Hspa2* (Dix et al., 1996; Dix et al., 1997), *Maps* (Li et al., 2021). They are not only downregulated in *Zfp541^-/-^* testes but also identified as target genes by ZFP541 CUT&Tag, suggesting that ZFP541 is able to prompt meiosis by directly regulating the transcription of meiotic essential genes. It is worth noting that the expression of *Kctd19* is regulated by ZFP541 but not vice versa. These results indicate that ZFP541 play a dominant role by binding *Kctd19* and regulating meiotic essential gene expression.

Horisawa-Takada *et al*. (2021) reported that HDAC1/2-containing ZFP541 complex directly represses the transcription of a subset of critical genes (Horisawa-Takada et al., 2021). HDAC1/2 complex is conventionally thought of as a genome-wide transcriptional repressor via histone deacetylation (Hassig et al., 1997; Kadosh and Struhl, 1998; Lee et al., 2018; Xue et al., 1998). In recent years, there have been many studies that HDAC1/2 tend to localize to transcriptionally active loci including promoters, gene bodies and enhancers (Jacob et al., 2011; Mondal et al., 2020; Wang et al., 2009). HDAC1/2 activity and targeting to specific gene loci strongly depends on the complexes they are involved in (Kelly and Cowley, 2013; Millard et al., 2017). Our data demonstrate that ZFP541 containing HDAC1/2 complex is enriched in the promotor of meiotic and post-meiotic genes, and activates their expression. We speculate the specific transcriptional activating capacity for HDAC1/2 is dependent on ZFP541. As a matter of fact, it makes sense that HDACs appears in transcriptional activating genome loci because the chromatin structure needed to be re-established quickly after mRNA transcription in order to avoid unexpected expression of adjacent genes and protect genome from excessive exposure.

In summary, our study demonstrated that ZFP541 associates with KCTD19 and HDAC1/HDAC2 to form a meiotic regulating complex which is essential for pachytene progression and fertility in male mice. In particular, ZFP541 binds to KCTD19 and is also able to activate the expression of *Kctd19*, forming an auto-regulatory circuit to guarantee the meiotic progression and spermatogenesis.

## Materials and methods

### Generation of Zfp541^-/-^, Kctd19^-/-^, Zfp541-EGFP and Kctd19-EGFP mice

*Zfp541*^-/-^ and *Kctd19*^-/-^ mice were generated by co-microinjection of *in vitro* transcribed *Cas9* mRNA and the synthetic sgRNAs into C57BL/6 zygotes. *Zfp541-EGFP* and *Kctd19-EGFP* mice were generated by co-microinjection of *in vitro* transcribed *Cas9* mRNA, the synthetic sgRNA and the targeting *EGFP* vector into C57BL/6 zygotes. The targeting strategy, including the gRNA sequences, the knockout or knock-in alleles obtained, are depicted respectively. The sgRNAs were prepared using a MEGAshortscript T7 Transcription kit (AM1354; Ambion, Austin, TX, USA) according to the manufacturer’s instructions. The T7-Cas9 DNA was amplified by PCR using plasmid hCas9 as the template and *in vitro* transcribed using a T7 Ultra Kit (AM1345; Ambion, TX, US). *Cas9* mRNA was purified using an RNeasy Mini Kit (74104; Qiagen, Venlo, Netherlands, Germany). The *Cas9* mRNA and sgRNAs were diluted into injection buffer (0.25 mM EDTA, 10 mM TrisHCl, pH 7.4) and incubated for 10 min at 37°C before injection. The final concentration of *Cas9* was 80 ng/ml, and that of sgRNA was 20 ng/ml. The donor plasmid concentration was 10 ng/ml for the injection. For microinjection, the *Cas9* mRNA, sgRNAs and the targeting vector were introduced into cytoplasm and the larger male pronucleus of C57BL/6 mouse zygotes by an inverted microscope (OLYMPUS IX71) equipped with a micro-injector system (Eppendorf FemtoJet 4i). Embryos were transferred to pseudopregnant C57BL/6 female mice according to standard procedures (20-30 zygotes per pseudopregnant mice). The primers used for genotyping are listed (*Supplementary file 3*). All animal studies were approved by the Chinese Ministry of Health national guidelines, and performed following institutional regulations after review and approval by the Institutional Animal Care and Use Committee at the National Institute of Biological Sciences, Beijing.

### Generation of anti-ZFP541, anti-KCTD19 and anti-H1t antibodies

Polyclonal rabbit antibodies against ZFP541 and H1t were produced by immunization with mouse polypeptides of ZFP541 (residues 32-46: C-TLNRDLGPSTRDLLY-NH2) (ChinaPeptides, Beijing, China) and mouse polypeptides of H1t (residues 18-32: Ac EKPSSKRRGKKPGLC) (Drabent et al., 1996) (ChinaPeptides). KCTD19 recombinant antibody was performed as previously described (Choi et al., 2008). All antibodies were generated and purified in the National institute of Biological Sciences Beijing, Antibody Centre.

### PCR, quantitative real-time PCR (qRT-PCR)

Genomic DNA was isolated from tail tip following the HotSHOT method (Truett et al., 2000) and genotyping was performed using standard PCR methods with sequence-specific primers. Total RNA was extracted from indicated tissues using Trizol reagent (15596026; Invitrogen, Carlsbad, CA, USA) following standard protocols, and was reverse transcribed using a PrimeScript ® RT reagent kit with gDNA Eraser (RR047A; TaKaRa, Dalian, China). qRT-PCR was performed with SYBR Green master mix (DRR420A; TaKaRa) using an ABI PRISM 7500 Sequence Detection System (Applied Biosystems). Relative mRNA expression levels were calculated using the comparative CT method (normalized to the expression of *Gapdh*). The primers used are listed in the Supplemental Materials (*Supplementary file 3*).

### RNA-Seq and data analysis

We performed RNA-seq on whole testes dissected away from P14 *Zfp541*^+/-^ and *Zfp541^-/-^* males. Each genotype was represented by three biological replicates of one pair of testis each. Total RNA (∼1 μg) were submitted to the Sequencing Center at National Institute of Biological Sciences for RNA-Seq library preparation, and sequencing. The libraries were prepared using standard Illumina protocol and sequenced on the Illumina HiSeq 2500 platform using the Single-End 1x75bp sequencing strategy. For analysis, raw transcriptome sequence data were aligned with the mouse genome (GRCm38/mm10) using HISATS2 (v2.1.0). HTseq-count (v0.11.0) was used to quantify the reads counts for each sample. DEseq2 (v3.8) was used to analyze fold changes of gene expression between *Zfp541*^+/-^ and *Zfp541^-/-^* samples. Genes with significantly different expression levels (mean RPKM ≥ 1, false discovery rate (FDR) < 0.05, fold change > 1.5, removing no annotation genes) were chosen for enrichment analysis. Gene ontology analysis and data visualization were done using Metascape.

### Histological and TUNEL analyses

Testes and caudal epididymides were freshly fixed in 4% paraformaldehyde (PFA) (DF0135; Beijing leagene biotech, Beijing, China) overnight at 4°C, dehydrated in an ethanol series (70%, 80%, 90%, 100%), and embedded in paraffin. 5 µm thickness sections were prepared and mounted on glass slides. After deparaffinization and re-hydration, the slides were stained with hematoxylin and eosin. For TUNEL analysis, assays were carried out using the in Situ Cell Death Detection Kit (11684795910; Roche, Basel, Switzerland) as previously described (Liu et al., 2016). The paraffin sections were treated with the TUNEL reaction mixture, and incubated in a humidified atmosphere for 60 min at 37°C in the dark.

### Immunofluorescence, chromosome spreads and antibodies

For paraffin section, the testes were fixed in 4% PFA in PBS and embedded in paraffin. Serial sections were dewaxed and rehydrated, and antigen retrieval was performed by microwaving the sections in 0.01 M sodium citrate buffer (pH 6.0). Sections were then blocked (10%FBS, 1% Triton X-100 in PBS) for 1 hour at room temperature, and incubated with primary antibodies overnight at 4°C. Subsequently, sections were washed and incubated with appropriate secondary antibodies with Alexa Fluor 488, 555 or 633 at 37°C for 1 hour, and counterstained with DAPI (1:1000; D1306; Invitrogen) for 10 min, and mounted on plus-coated slides that were cover-slipped using vectashield antifade mounting medium (H-1000; Vector Laboratories, Burlingame, CA, USA). Finally, sections were photographed using Nikon SIM confocol microscope.

For frozen section, testes were fixed in 4% PFA for overnight at 4°C, and then dehydrated in 30% sucrose/PBS. Tissue was embedded in Cryo-gel Tissue-Tek OCT compound (62806-01; Tissue-Tek, Torrance, CA, USA) and frozen on dry ice. Cryosections of 10 µm thickness were cut from snap-frozen samples using a cryomicrotome (CM1950; Leica Biosystems, Wetzlar, Germany). Cryosectioned samples were used for tissue staining and stored at -20°C. Chromosome spreads of spermatocytes were performed as previously described (Peters et al., 1997) and were further staining.

Primary antibody used were rabbit antibody to SYCP3 (1:500; NB300-232; Novus, CO, USA), mouse antibody to SYCP3 (1:100; sc-74569; Santa Cruz Biotechnology, CA, USA), goat antibody to SYCP3 (1:100; sc-20845; Santa Cruz Biotechnology), rabbit antibody to SYCP1 (1:500; NB300-229; Novus), mouse antibody to γH2AX (1:500; 05-636; MilliporeSigma, St. Louis, MO), mouse antibody to DMC1 (1:100; sc-22768; Santa Cruz Biotechnology), mouse antibody to MLH1 (1:50; 550838; BD Pharmingen™, San Diego, CA, USA), rabbit antibody to H3K9me3 (1:500; 39161; Active Motif, Palo Alto, CA, USA), rabbit antibody to H3K9ac (1:1000; ab177177; Abcam, MA, USA), chicken antibody to GFP (1:500; ab13970; Abcam), mouse antibody to α-TUBULIN (1:500; T6199-100U; MilliporeSigma), human antibody to CREST(1:500; HCT-0100; Immunovision, Springdale), mouse antibody to RNA polymerase II (1:100; sc-47701; Santa Cruz Biotechnology). Secondary antibodies used were 488 conjugated goat anti-chicken IgG (1:500; A-11039; Invitrogen), 488 conjugated donkey anti-mouse IgG (1:500; A-21202; Invitrogen), 488 conjugated donkey anti-rabbit IgG (1:500; A-31571; Invitrogen), 555 conjugated donkey anti-mouse IgG (1:500; A-31570; Invitrogen), 555 conjugated donkey anti-rabbit IgG (1:500; A-31572; Invitrogen), 633 conjugated goat Anti-rabbit IgG (1:500; A-21070; Invitrogen).

### Immunohistochemistry

Immunohistochemistry was performed using a SP link Detection Kit (Biotin-Streptavidin HRP Detection Systems) (SP-9001; ZSGB-Bio, Beijing, China). Briefly, sections were then blocked using normal goat serum for 30 min at room temperature, and incubated with primary antibodies (ZFP541 and KCTD19, homemade) overnight at 4°C. Sections were washed and incubated with a biotinylated goat anti-rabbit IgG for 15 min at room temperature. Slides were then washed and incubated for horseradish peroxidase-conjugated streptavidin for 15 min at room temperature. Peroxidase activity was detected with a diaminobenzidine (DAB) Peroxidase Substrate Kit (ZLI9017; ZSGB-Bio). Sections were counter-stained with hematoxylin, dehydrated, and covered with glass coverslips.

### Vector construction and cell transfections

The full-length coding sequences of mouse *Zfp541*, *Kctd19*, *Hdac1*, *Hdac2* and *Dnttip* were amplified from mouse testis cDNA by PCR. Primers are listed (*Supplementary file 3*). These fragments were cloned into pcDNA3.1-N-3XFLAG, pcDNA3.1/myc-His B or pEGFP-N1 plasmid to synthesize recombinant protein. Each required tagged protein expression construct was transfected in HEK 293T cells with Lipofectamine 3000 transfection reagent (L3000150; Invitrogen) according to the manufacturer’s protocol. Cells were collected 48 hours after transfection, performed by Co-IP with anti-Flag antibody (F1804; MilliporeSigma), and followed by western blotting analysis with anti-Flag and anti-Myc antibodies (16286-1-AP; Proteintech, Wuhan, China). For immunostaining: Cells were transfected with the ZFP541-pEGFP-N1 and MYC-tagged expression constructs at 48 hours after transfection, and immune-stained for GFP and MYC.

### Western blotting

Testes were rinsed with PBS and lysed in cold RIPA buffer (9806; Cell Signaling, Danvers, MA, USA) supplemented with complete protease inhibitor cocktail (11697498001; Roche). Homogenized lysates were rotated for 1 hour at 4°C, and centrifuged at 13000×g for 20 min at 4°C.The protein concentrate on of collected supernatant was determined using bicinchoninic acid (BCA) assay (23225; Thermo Fisher Scientific, Rockford, IL, USA). An equal amount of each protein sample was electrophoresed on sodium dodecyl sulfate-polyacrylamide gel electrophoresis (SDS-PAGE) and transferred onto polyvinylidene difluoride (PVDF) membranes (3010040001; Roche). The membrane was blocked with 5% non-fat dry milk for 2 hours at room temperature, and respectively incubated with primary antibodies at 4°C overnight. Primary antibody dilution: ZFP541 and KCTD19 (1:2000; homemade), HDAC1 (1:5000; ab19845; Abcam), HDAC2 (1:5000; ab12169; Abcam), DNTTIP1 (1:1000; A15558, ABclonal, Wuhan, China), FLAG (1:2000; F1804; MilliporeSigma), MYC (1:1000; 16286-1-AP; Proteintech). The PVDF membrane was then washed three times in 0.1% Tween-20 in Tris-buffered saline (TBST) and incubated with horseradish peroxidase-conjugated goat anti-rabbit IgG (1:5000; A6154; MilliporeSigma) or goat anti-mouse IgG (1:5000; A4416; MilliporeSigma) for 1 hour at room temperature. After washing with TBST, the membrane was treated with the Pierce™ ECL 2 western blotting Substrate (34577; Thermo Fisher Scientific).

### Immunoprecipitation and Mass spectrometry (MS)

Testes (50 mg) or HEK 293T cells (1x10^7^) were homogenized in 500 μl ice-cold lysis buffer (50 mM Tris, pH 7.6, 150 mM NaCl, 2 mM EDTA, 1% Triton X-100, 10% glycerol, 1x protease inhibitor) on ice for 15 min. The testis lysates were centrifuged at 13000×g for 20 min at 4°C. The supernatants were collected and incubated with 30 µl Dynabeads Protein A (10002D; Invitrogen) or Dynabeads Protein G (10004D; Invitrogen) and 8 µg indicated antibodies overnight at 4°C with rotation. The samples were applied to columns in the magnetic field of the Millipore separator, and the columns were washed four times with lysis buffer. Finally, protein binding on the beads were eluted with elution buffer (pH=3, 1M glycine).

For ZFP541 IP-MS, immunoprecipitations were divided into wild-type and ZPF541-EGFP lysate with GFP antibody using P18 mouse testes. For KCTD19 IP-MS, immunoprecipitations were divided into wild-type and *Kctd19^-/-^* lysate with KCTD19 antibody using P18 mouse testes. The immunoprecipitated proteins were subjected to SDS-PAGE followed by silver staining (PROT-SIL1; MilliporeSigma). The silver-stained proteins were destained and digested in-gel with trypsin at 37°C overnight. The digested peptides were eluted on a capillary column and sprayed into a Q Exactive Mass Spectrometer (Thermo Fisher Scientific) equipped with a nano-ESI ion source (National institute of Bilogical Sciences, Beijing, Proteomics Center).

### Isolation of spermatogenic cells and Flow cytometry purification of spermatocytes

Mouse testicular cell suspensions were isolated from *Zfp541^+/-^* and *Zfp541^-/-^* testes as described (Chen et al., 2018; Gaysinskaya et al., 2014). Briefly, the seminiferous tubules were minced and digested at a final concentration of 1 mg/ml collagenase IV and 1mg/ml DNase I at 37°C for 20 min, settled by standing the tube vertically, and washed with DMEM. Cell suspension was subsequently digested with 0.25% Trypsin containing 1mg/ml DNase I 37^◦^C for 5 min to prepare single-cell suspensions. Enzymatic digestion was quenched with DMEM supplemented with 10% FBS. Single cell suspensions were sieved through a 40 μm cell strainer, and stained with Hoechst 33342 (B2261; MilliporeSigma) at a final concentration of 10 µg/ml for 1 hour at 37°C. Prior of sorting, propidium iodide (P3566; Invitrogen) was added to a final concentration of 2 µg/ml. Cell suspensions were sorted on an BD FACSAria Fusion-II (Becton Dickinson) using a 70 µm nozzle. Hoechst was excited with a UV laser at 355 nm and fluorescence was recorded with a 450/40 nm band-pass filter (Hoechst blue) and 635 nm long filter (Hoechst red). Spermatocytes were sorted and collected in DMEM containing 10% FBS.

### CUT&Tag and data analysis

CUT&Tag assay was performed as described previously with modifications (Kaya-Okur et al., 2019). 1×10^5^ testicular single cells were washed twice in 1 ml of Wash Buffer (20 mM HEPES pH 7.5, 150 mM NaCl, 0.5 mM Spermidine, 1× Protease inhibitor cocktail) and centrifuged at 400 g for 5 min. Concanavalin A-coated magnetic beads (BP531; Bangs Laboratories, IN, USA) were washed twice with binding buffer (20 mM HEPES pH 7.5, 10 mM KCl, 1 mM MnCl_2_, 1 mM CaCl_2_). Next, 10 μl of activated beads were added per sample and incubated at room temperature for 15 min. The unbound supernatant was removed and bead-bound cells were resuspended in 50 μl of Dig-wash buffer (20 mM HEPES pH 7.5, 150 mM NaCl, 0.5 mM Spermidine, 0.05% Digitonin, 1× Protease inhibitor cocktail) containing 2 mM EDTA, 0.1% BSA and 1 μg rabbit polyclonal GFP antibody (ab290; Abcam). The GFP antibody was incubated on a rotating platform overnight at 4°C, and removed using a magnet stand. Guinea Pig anti-Rabbit IgG (Heavy & Light Chain) secondary antibody (ABIN101961; antibodies-online) was diluted in 100 µl of Dig-wash buffer and cells were incubated for 1 hour. Cells were washed three times with Dig-wash buffer to remove unbound antibody. The Hyperactive pA-Tn5 Transposase adapter complex (TTE mix) (S603-01; Vazyme, Nanjing, China) was diluted 1:100 (0.04 μM) in Dig-med buffer (20 mM HEPES pH 7.5, 300 mM NaCl, 0.5mM Spermidine, 0.01% Digitonin, 1× Protease inhibitor cocktail). Cells were incubated with 0.04 μM TTE mix at room temperature for 1 hour, washed three times with Dig-med buffer to remove unbound pA-Tn5 protein. Cells were then resuspended in 300 μl of Tagmentation buffer (10 mM MgCl_2_ in Dig-med buffer) and incubated at 37°C for 1 hour. To terminate tagmentation, 10 μl of 0.5 M EDTA, 3 μl of 10% SDS and 5 μl of 10 mg/ml Proteinase K were added to 300 μl of per sample and incubated at 55 °C for 1 hour. DNA was purified using phenol-chloroform-isoamyl alcohol extraction and ethanol precipitation as well as RNase A treatment.

For library amplification, 24 μl of DNA was mixed with 10 μl of 5×Multiplex PCR Mix (E086-YSAA; Novoprotein, NJ, USA),14 μl of ddH2O, as well as 1 μl of uniquely barcoded i5 and i7 primers. A total volume of 50 μl of sample was placed in a PCR amplifier. To purify the PCR products, 1.2× volumes of VAHTS DNA Clean Beads (N411-02; Vazyme) were added and incubated at room temperature for 10 min. Libraries were washed twice with 80% ethanol and eluted in 22 µl of ddH2O. Libraries were sequenced on an Illumina NovaSeq platform and 150-bp paired-end reads were generated (Novogene, Beijing, China).

CUT&Tag paired reads were trimmed to remove adapter sequence using cutadapt. The trimmed reads were mapped to the UCSC mm10 genome assemblies using Bowtie2 v2.2.5 with very-sensitive parameters. Genome-wide normalized signal coverage tracks were created by bamCoverage in deepTools (version 3.4.3) and visualized in the Integrative Genomics Viewer (IGV version 2.4.9). Peak calling was carried out using MACS program with default parameters (version 2.2.7.1). Peaks were annotated to the genomic region and the nearest genes within 2 kb of TSS using Bioconductor package ChIPSeeker (version 1.22.1). Peaks over lapping by at least 1nt with unique gene model promoters (±3 kb of each unique gene model Transcription Starting Site) were considered as promoter located. *De novo* motif identification of CUT&Tag peaks were performed using Homer (version 4.11.1) website with default parameters (http://homer.ucsd.edu/homer/motif/).

### Statistical analysis

All data are reported as the means ± SD. Significance was tested by using the two-tailed unpaired Student’s *t*-test (*\*P* < 0.05; *\*\*P* < 0.01; *\*\*\* P* < 0.001) using GraphPad Prism 6 (GraphPad Software).

## Acknowledgements

We would like to thank Yanna Sun and Yue Yin for help with the RNA-seq experiment. We would like to thank Jinmei Chen for assistance with flow cytometry purification of mouse spermatocytes. We thank Hui Han for mass spectrometry and data analysis. This work was supported by the National Natural Science Foundation of China (81901537, YSL); Key Technologies Research and Development Program of Henan Province (192102310131, YSL); and Xinxiang Medical University (XYBSKYZZ201802, YSL).

## Author contributions

Yushan Li, Formal analysis, Supervision, Funding acquisition, Investigation, Methodology, Writing-original draft, Writing-review and editing; Ranran Meng, Shanze Li, Formal analysis, Investigation, Methodology; Bowen Gu, Methodology, Writing-review and editing; Xiaotong Xu, Haihang Zhang, Tianyu Shao, Jiawen Wang, Yinghua Zhuang, Methodology; Fengchao Wang, Conceptualization, Formal analysis, Supervision, Funding acquisition, Investigation, Methodology, Writing-review and editing

---

**Figure 1-figure supplement 1.**
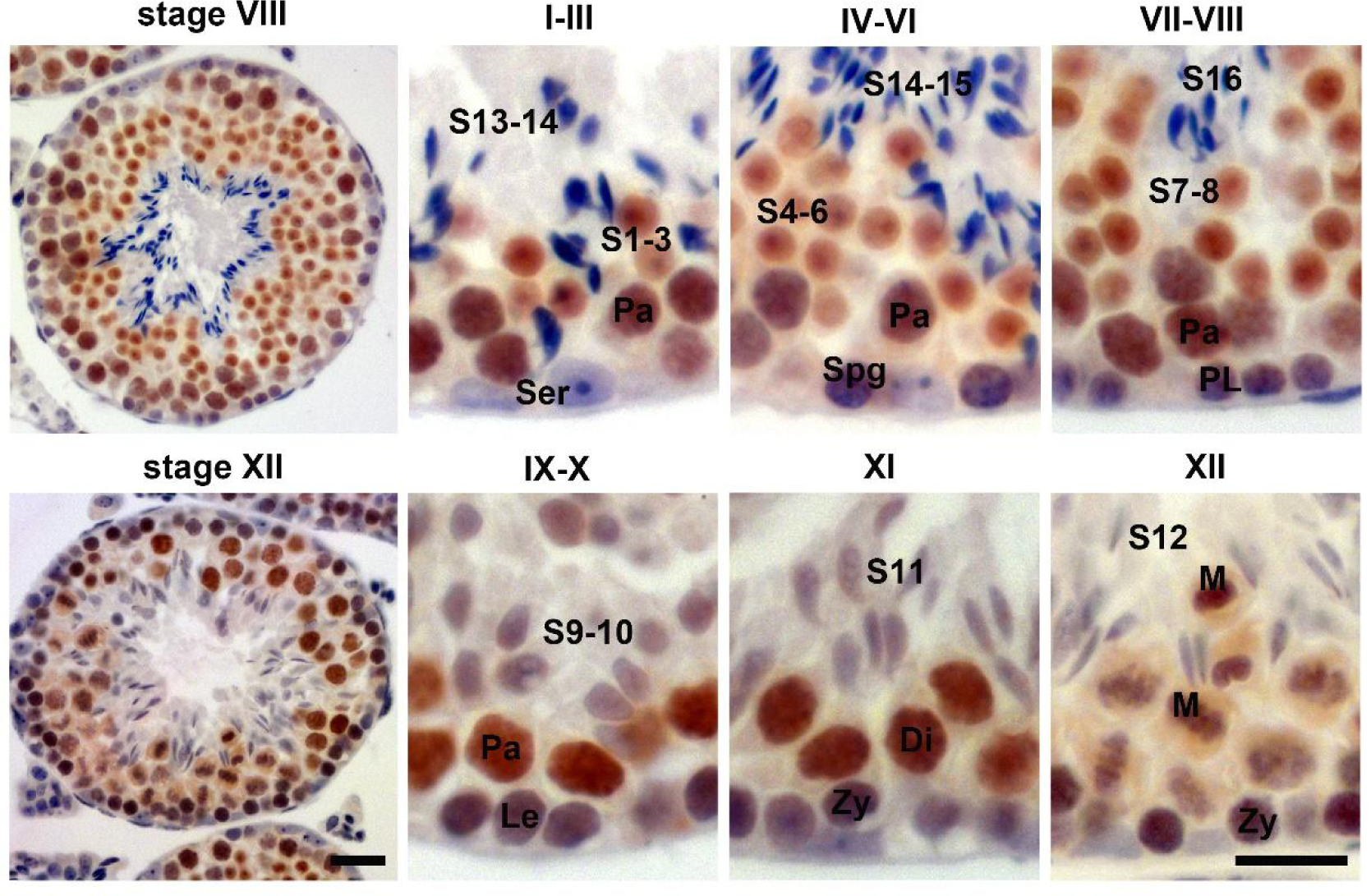
ZFP541 is expressed from leptotene spermatocytes to round spermatids in male mice. Immunohistochemistry analysis of ZFP541 in the 10-week-old mouse testes. Sertoli cells (Ser), Spermatogonia (Spg), Preleptotene (PL), Leptotene (Le), Zygotene (Zy), Pachytene (Pa), Diplotene (Di), Metaphase (M), spermatids (S). Scale bars: 50 μm.

**Figure 1-figure supplement 2.**
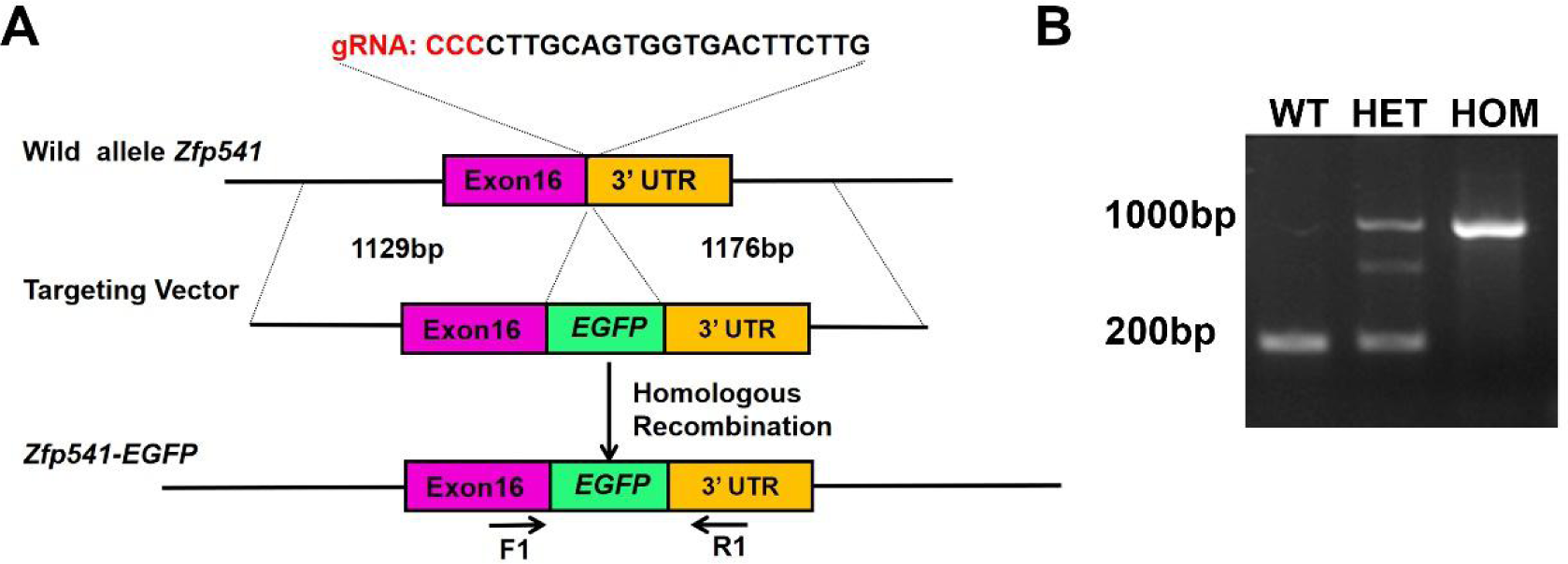
Generation of *Zfp541* C-terminus *EGFP* (*Zfp541-EGFP)* knock-in mice. **(A)** Schematic overview of CRISPR/Cas9-mediated knock-in of the EGFP cassette at the *Zfp541* locus. The top panel shows the *Zfp541* genomic locus. The middle panel shows the design of the *Zfp541-EGFP* targeting donor. The bottom panel shows the design for how the targeting donor is recombined into the *Zfp541* genomic locus via CRISPR/Cas9-mediated homologous recombination. The locations of gRNA and primers (F1 and R1) are indicated. **(B)** Genotyping of mouse tail tip DNA by PCR amplification with primers (F1 and R1) indicated in (A). The PCR products size is indicated on the image.

**Figure 2-figure supplement 1.**
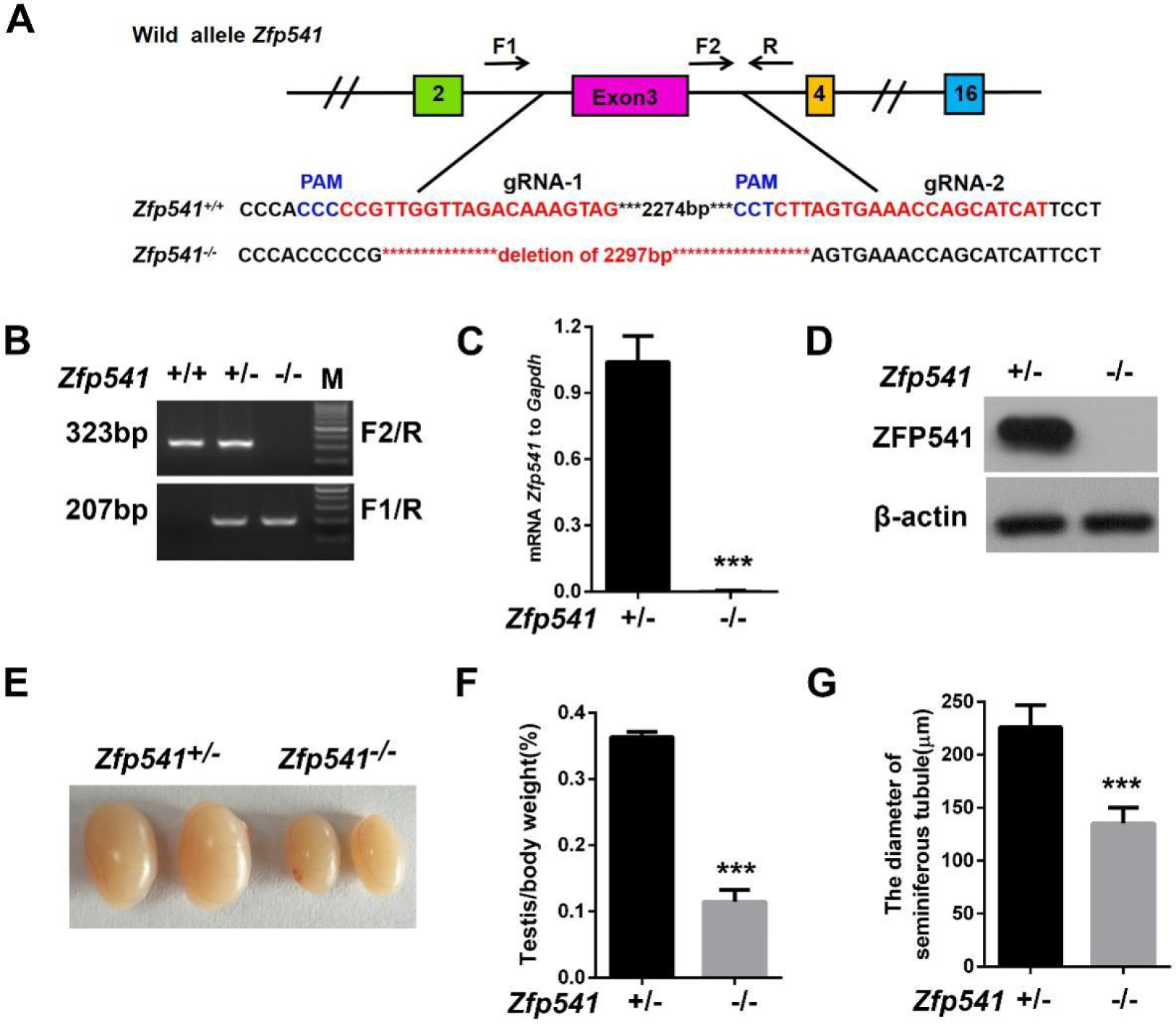
*Zfp541* is required for male fertility. **(A)** Schematic diagram showing the gene structure of *Zfp541* and the CRISPR/Cas9 strategy used to generate the knockout allele. The upper panel shows the *Zfp541* locus region to be targeted. Two gRNAs matching the exon 3 from its two ends were used to achieve deletion of a genomic fragment containing exon 3. The shift mutated sequences of the knockout allele are in the lower panel. The locations of gRNA and primers (F1, F2 and R) are indicated. **(B)** Genotyping of mouse DNA by PCR amplification with primers indicated in (A), the size of PCR products is indicated on the image. **(C)** qRT-PCR analysis of *Zfp541* mRNA level in 10-week-old testes of *Zfp541*^+/-^ and *Zfp541*^-/-^ males, all results were normalized to levels of *Gapdh* (n = 3, *\*\*\*P* < 0.001, Student’s *t*-test). **(D)** Western blotting analysis of ZFP541 protein level in testes between 10-week-old *Zfp541*^+/-^ and *Zfp541*^-/-^ males. β-actin was used as a loading control. **(E)** Representative morphology of testes of 10-week-old *Zfp541*^+/-^ and *Zfp541*^-/-^ males. **(F-G)** The testis to body weight ratio, and the diameter of the seminiferous tubules between 10-week-old *Zfp541*^+/-^ and *Zfp541*^-/-^ males (n = 4, *\*\*\*P* < 0.001, Student’s *t*-test).

**Figure 3-figure supplement 1.**
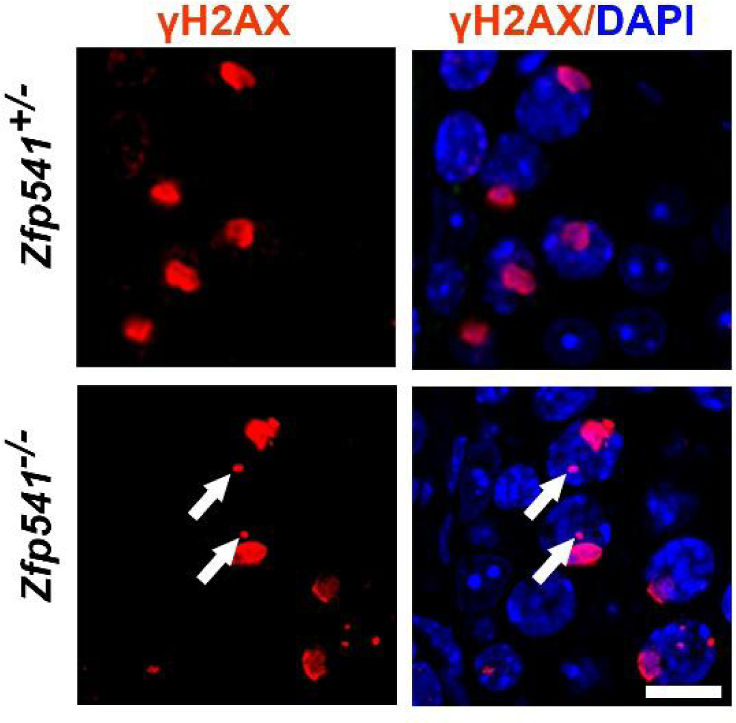
DSB repair is impaired in the *Zfp541^-/-^* spermatocytes. Immunofluorescence staining of γH2AX in paraffin sections of 10-week-old *Zfp541*^+/−^ and *Zfp541*^−/−^ testes. Arrows indicate abnormal γH2AX signals. Scale bars: 10 μm.

**Figure 3-figure supplement 2.**
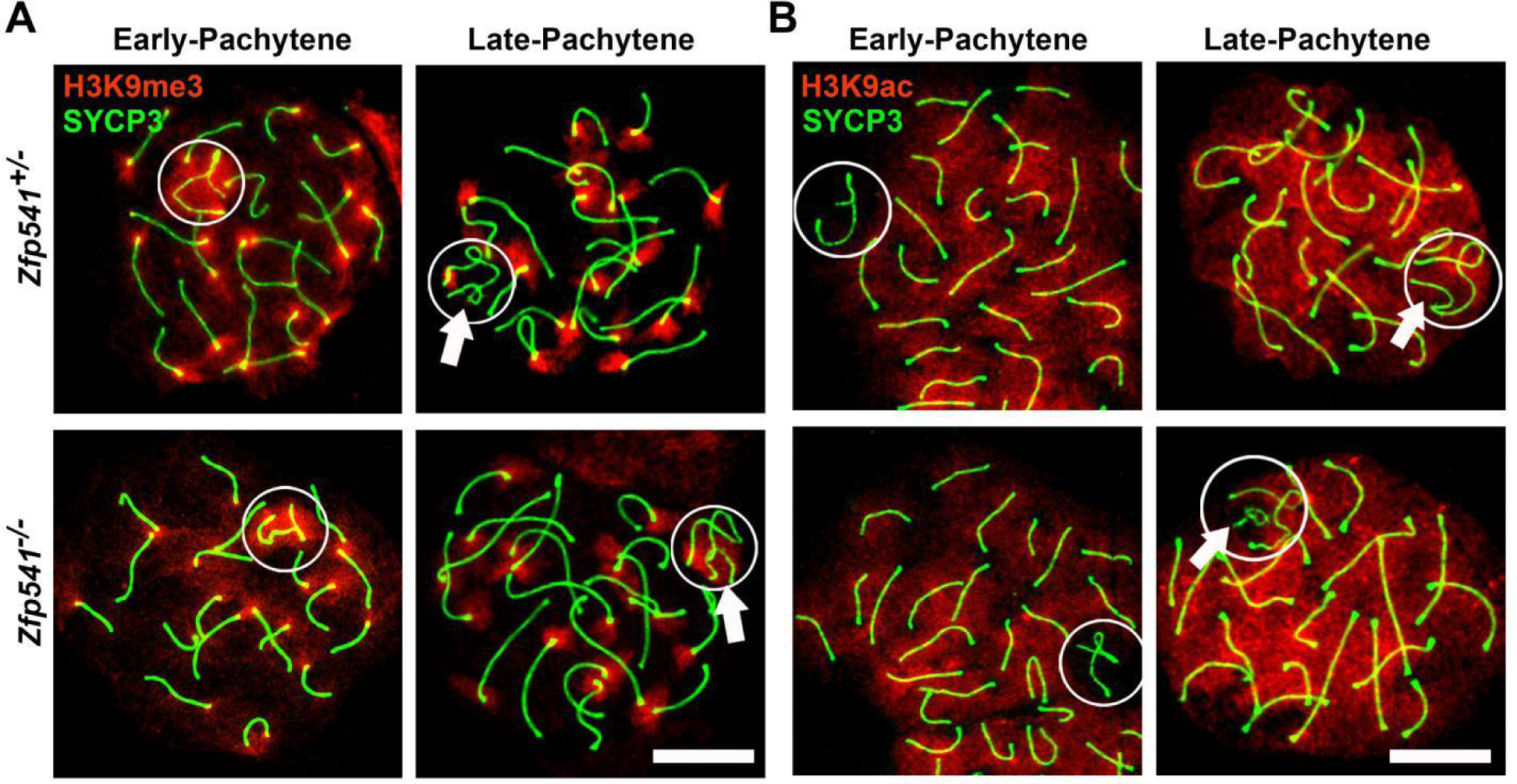
Depletion of *Zfp541* affects the localization of H3K9me3 and H3K9ac in the Y chromosome. **(A)** Immunofluorescent staining of SYCP3 and H3K9me3 on chromosome spreads of *Zfp541*^+/−^ and *Zfp541*^−/−^ pachytene spermatocytes. Circles indicate XY bodies, and arrows indicate Y chromosomes. SYCP3 staining (green); H3K9me3 staining (red); Scale bars: 10 μm. **(B)** Immunofluorescent staining of SYCP3 and H3K9ac on chromosome spreads of *Zfp541*^+/−^ and *Zfp541*^−/−^ pachytene spermatocytes. Circles identify XY bodies and arrows identify Y chromosomes. SYCP3 staining (green); H3K9ac staining (red). Scale bars: 10 μm.

**Figure 3-figure supplement 3.**
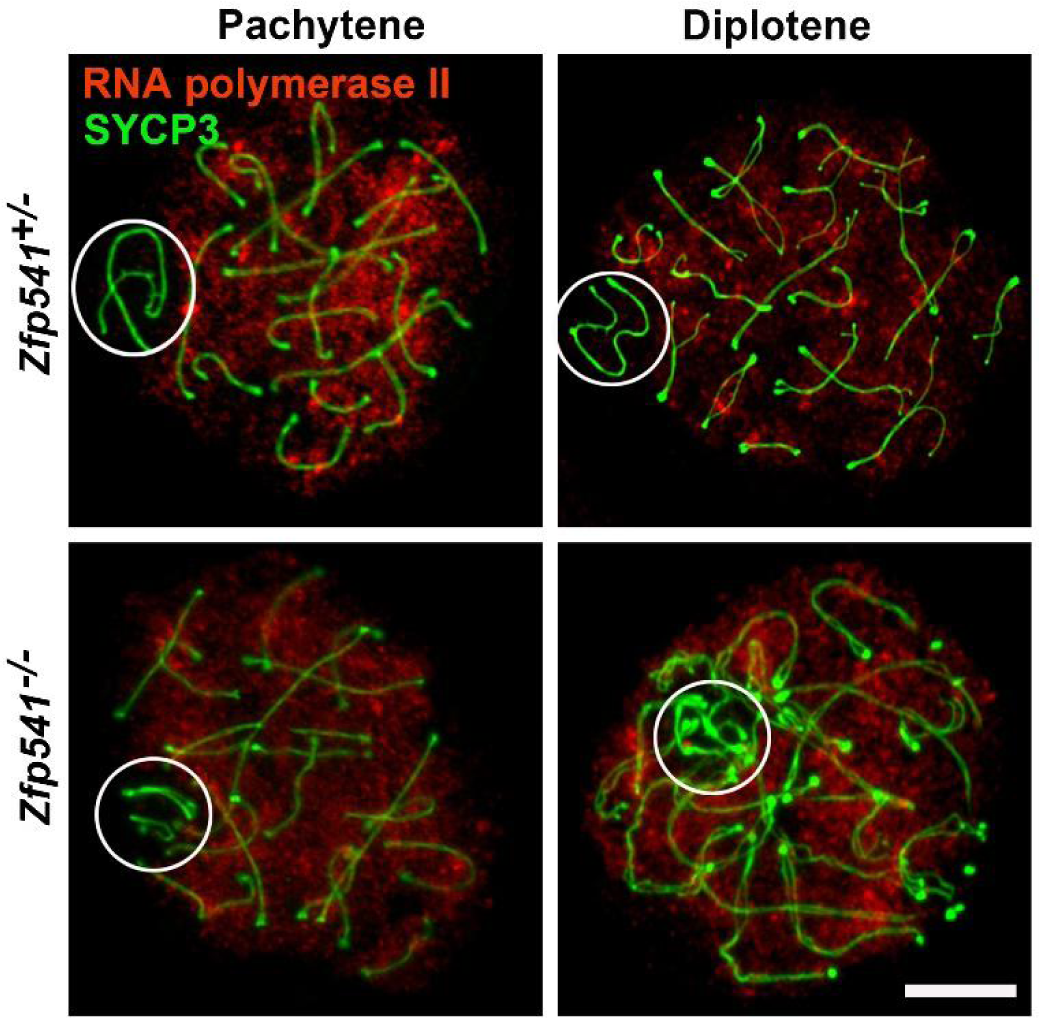
ZFP541 is dispensable for the maintenance of MSCI. Immunofluorescent staining of SYCP3 and RNA polymerase II on chromosome spreads of 10-week-old *Zfp541*^+/-^ and *Zfp541*^−/−^ pachytene and diplotene spermatocytes. SYCP3 staining (green); RNA polymerase II staining (red). Circles identify XY bodies. Scale bars: 10 μm.

**Figure 4-figure supplement 1.**
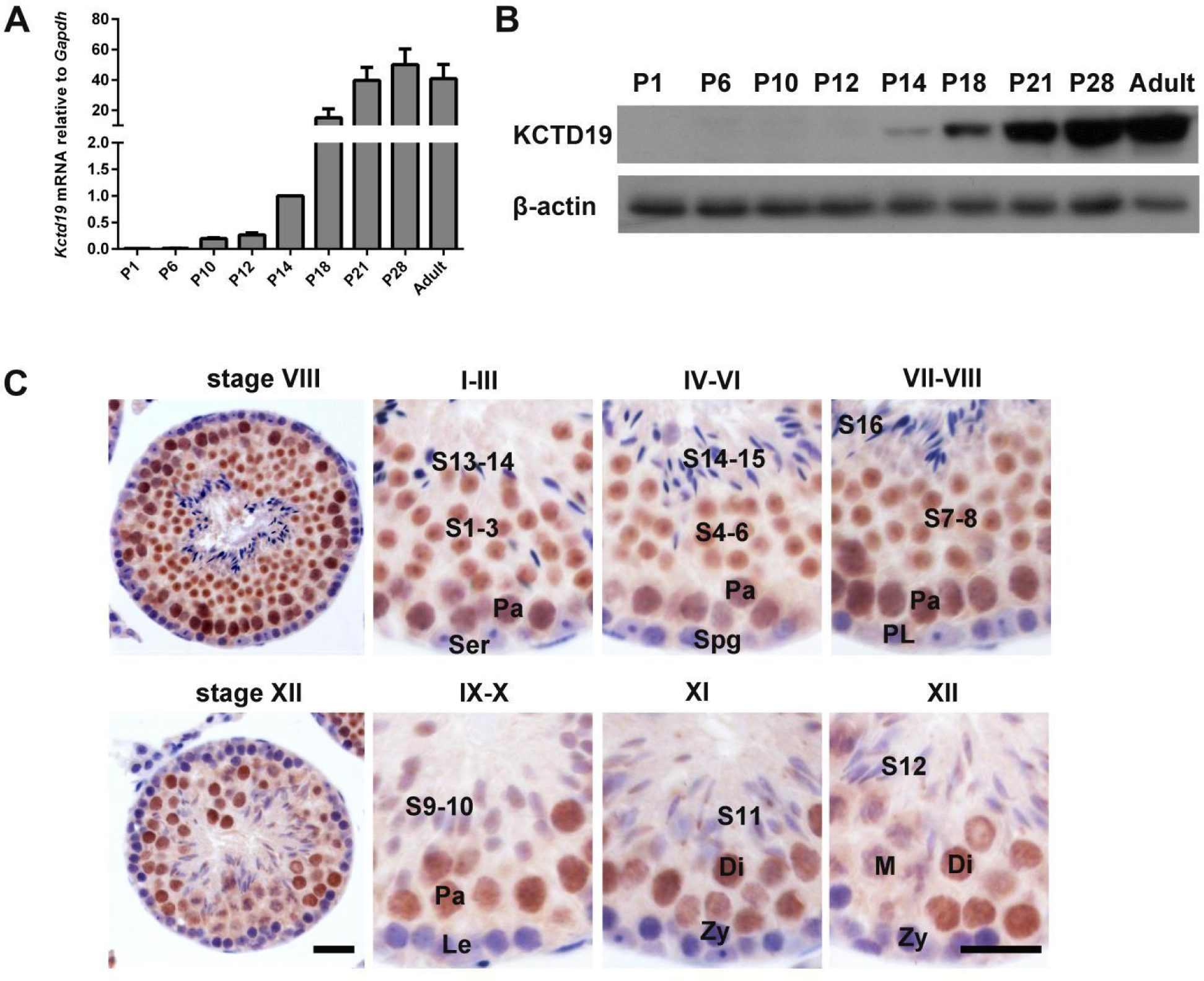
Expression pattern of *Kctd19* in mouse testes. **(A)** qRT-PCR analysis of the relative expression of *Kctd19* mRNA obtained from mouse testes collected at different time points during postnatal development, all data normalized to *Gapdh* mRNA levels (n = 3, Student’s *t*-test). **(B)** Western blotting analysis of KCTD19 protein obtained from mouse testes collected at different time points during postnatal development; β-actin was used as a loading control. **(C)** Immunohistochemical staining of KCTD19 protein in 10-week-old mouse testes. Sertoli cells (Ser), Spermatogonia (Spg), Preleptotene (PL), Leptotene (Le), Zygotene (Zy), Pachytene (Pa), Diplotene (Di), Metaphase (M), spermatids (S). Scale bars: 50 μm.

**Figure 4-figure supplement 2.**
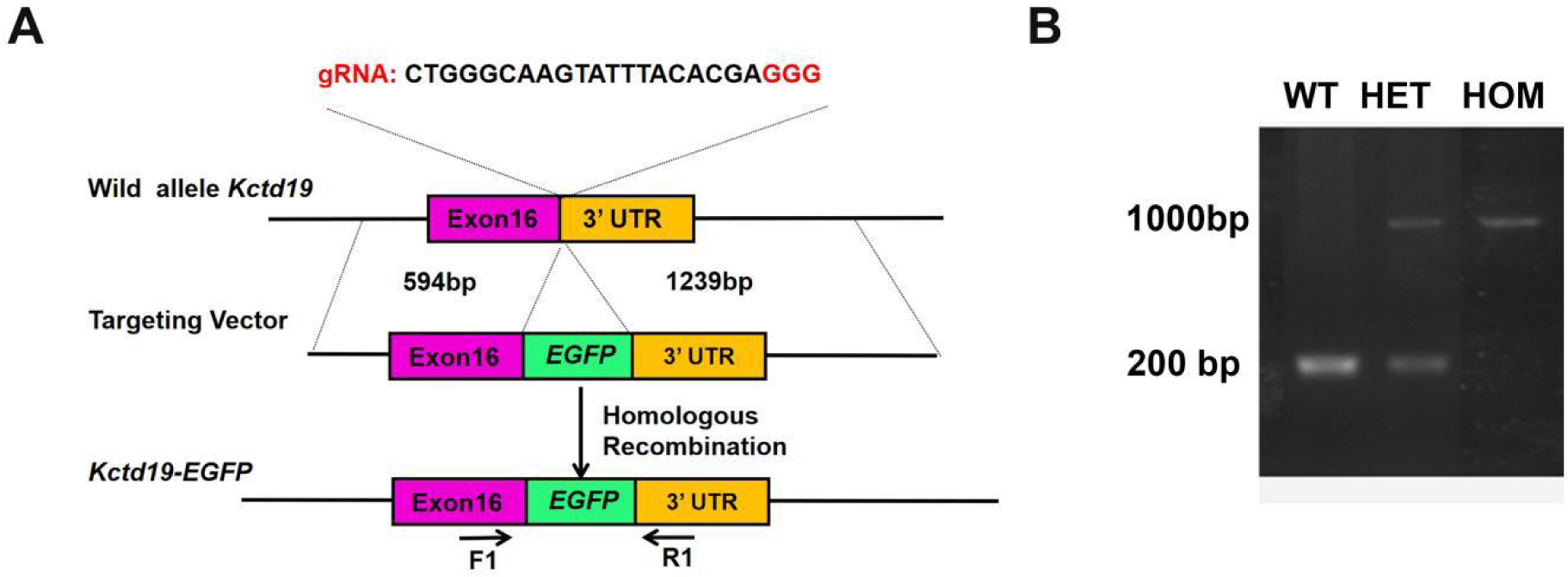
Generation of *Kctd19* C-terminus *EGFP* (*Kctd19-EGFP*) knock-in mice. **(A)** Schematic overview of CRISPR/Cas9-mediated knockin of the EGFP cassette at the *Kctd19* locus. The top panel shows the *Kctd19* genomic locus. The middle panel shows the design of the *Kctd19-EGFP* targeting donor. The bottom panel shows the design for how the targeting donor is recombined into the *Kctd19* genomic locus via CRISPR/Cas9-mediated homologous recombination in mice. The locations of gRNA and primers (F1 and R1) are indicated. **(B)** Genotyping of mouse tail tip DNA by PCR amplification with primers (F1 and R1) indicated in (A), the PCR product size is indicated.

**Figure 4-figure supplement 3.**
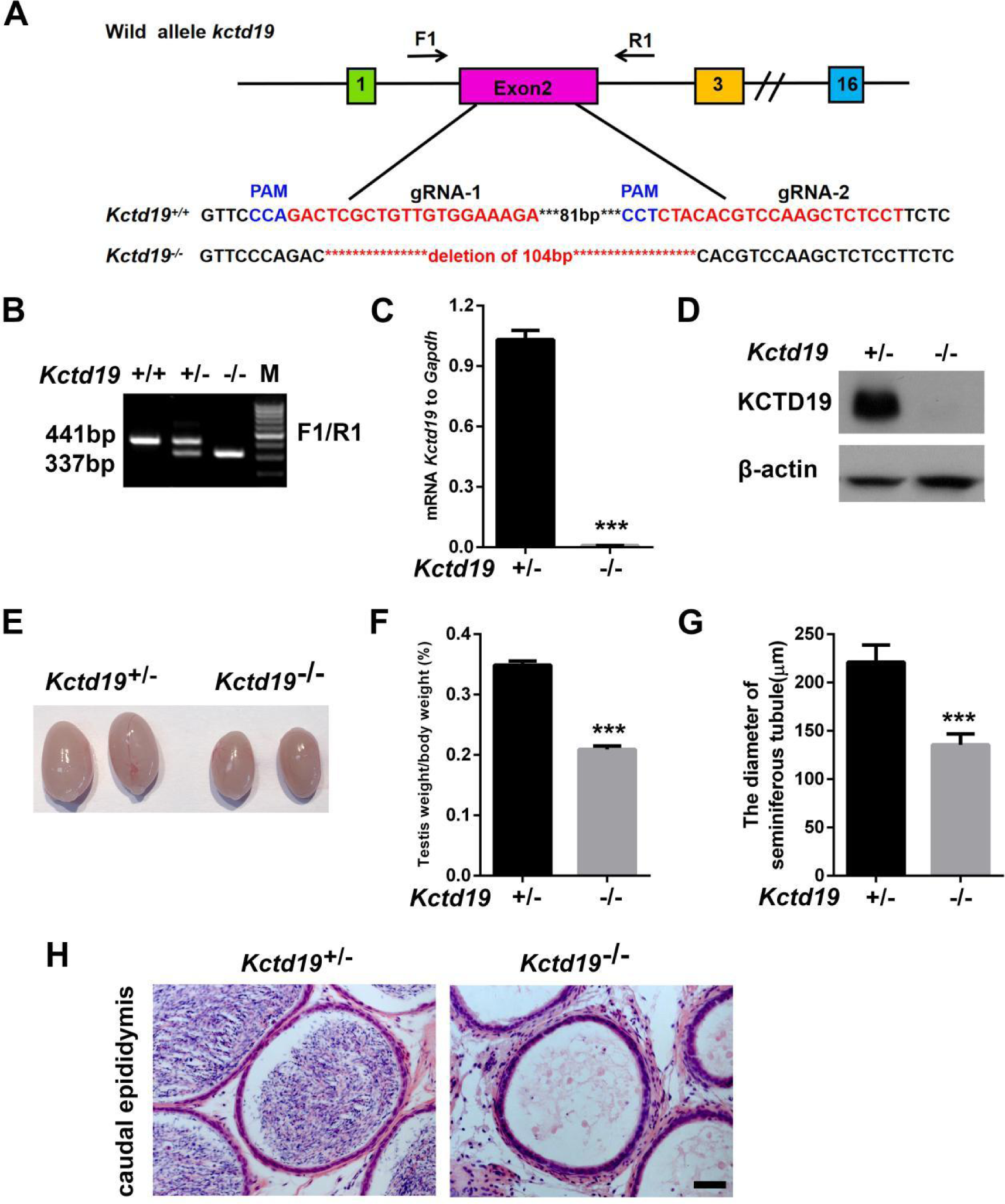
*Kctd19* is required for male fertility. **(A)** Schematic diagram showing the gene structure of *Kctd19* and the CRISPR/Cas9 strategy used to generate the knockout allele. The upper panel shows the targeted *Kctd19* locus region. Two gRNAs matching the exon 2 from interior were used to achieve deletion of a genomic fragment. The shift mutated sequences of the knockout allele are in the lower panel. The locations of gRNA and primers (F1 and R1) are indicated. **(B)** Genotyping of mouse tail tip DNA by PCR amplification with primers indicated in (A), the size of PCR products is indicated on the image. **(C)** qRT-PCR analysis of *Kctd19* mRNA level in testes from 10-week-old *Kctd19*^+/-^ and *Kctd19*^-/-^ males, all results were normalized to levels of *Gapdh* (n = 4, *\*\*\*P* < 0.001, Student’s *t*-test). **(D)** Western blotting analysis of testes from 10-week-old *Kctd19*^+/-^ and *Kctd19*^-/-^ males. β-actin was used as a loading control. **(E)** Representative image analysis of the morphology of testes derived from 10-week-old *Kctd19*^+/-^ and *Kctd19*^-/-^ males. **(F-G)** The testis to body weight ratio and the diameter of the seminiferous tubules between 10-week-old *Kctd19*^+/-^ and *Kctd19*^-/-^ males (n = 4, *\*\*\*P* < 0.001, Student’s *t*-test). **(H)** Hematoxylin and eosin staining of caudal epididymides in 10-week-old *Kctd19*^+/-^ and *Kctd19*^−/−^ males. Scale bars: 50 μm.

**Figure 4-figure supplement 4.**
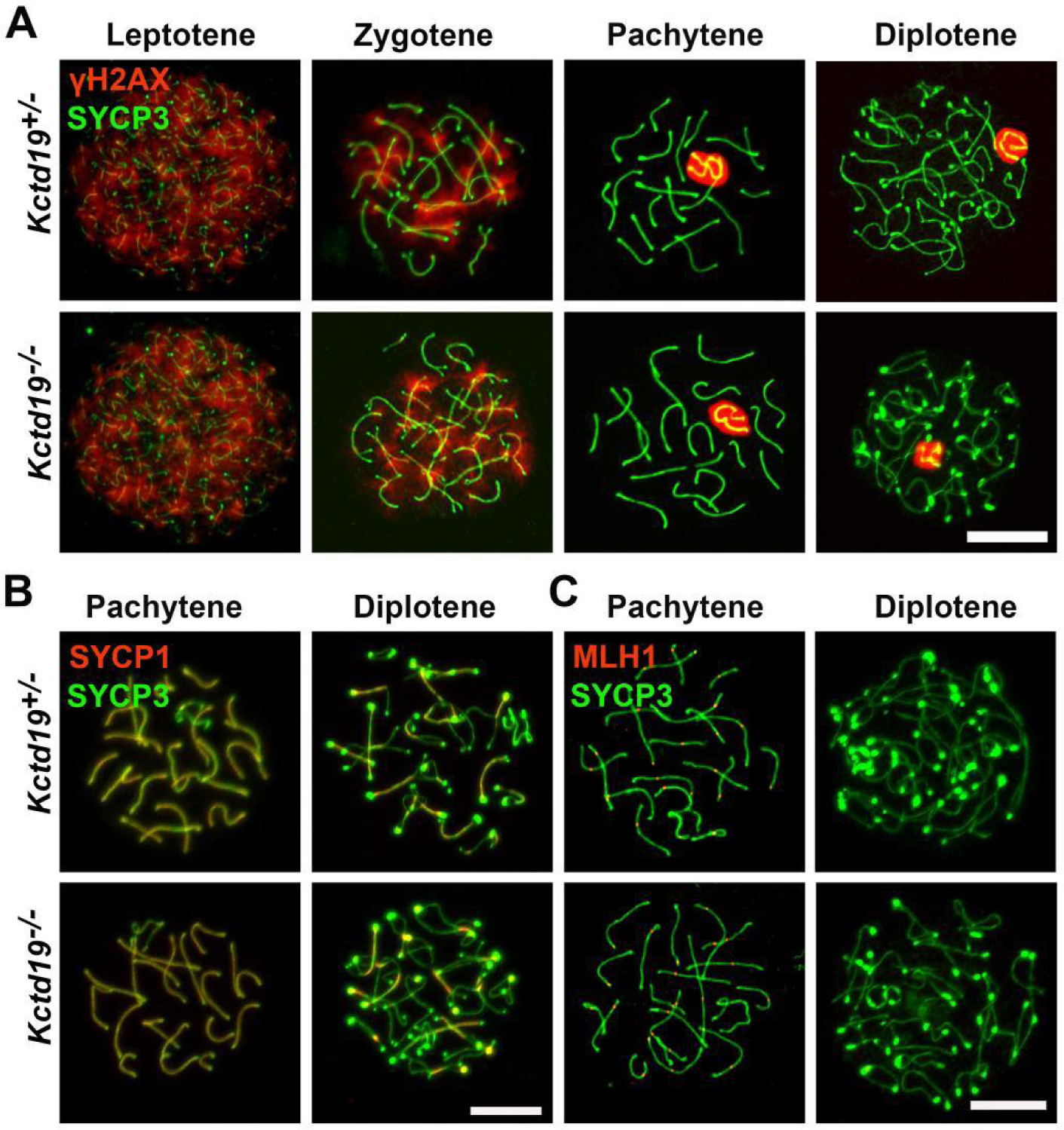
*Kctd19* is dispensable for homologous recombination. **(A-C)** Immunofluorescent staining of SYCP3, γH2AX, SYCP1, and MLH1 on chromosome spreads of 10-week-old *Kctd19*^+/-^ and *Kctd19*^−/−^ spermatocytes. SYCP3 staining (green); γH2AX, SYCP1, MLH1 staining (red). Scale bars: 10 μm.

**Figure 4-figure supplement 5.**
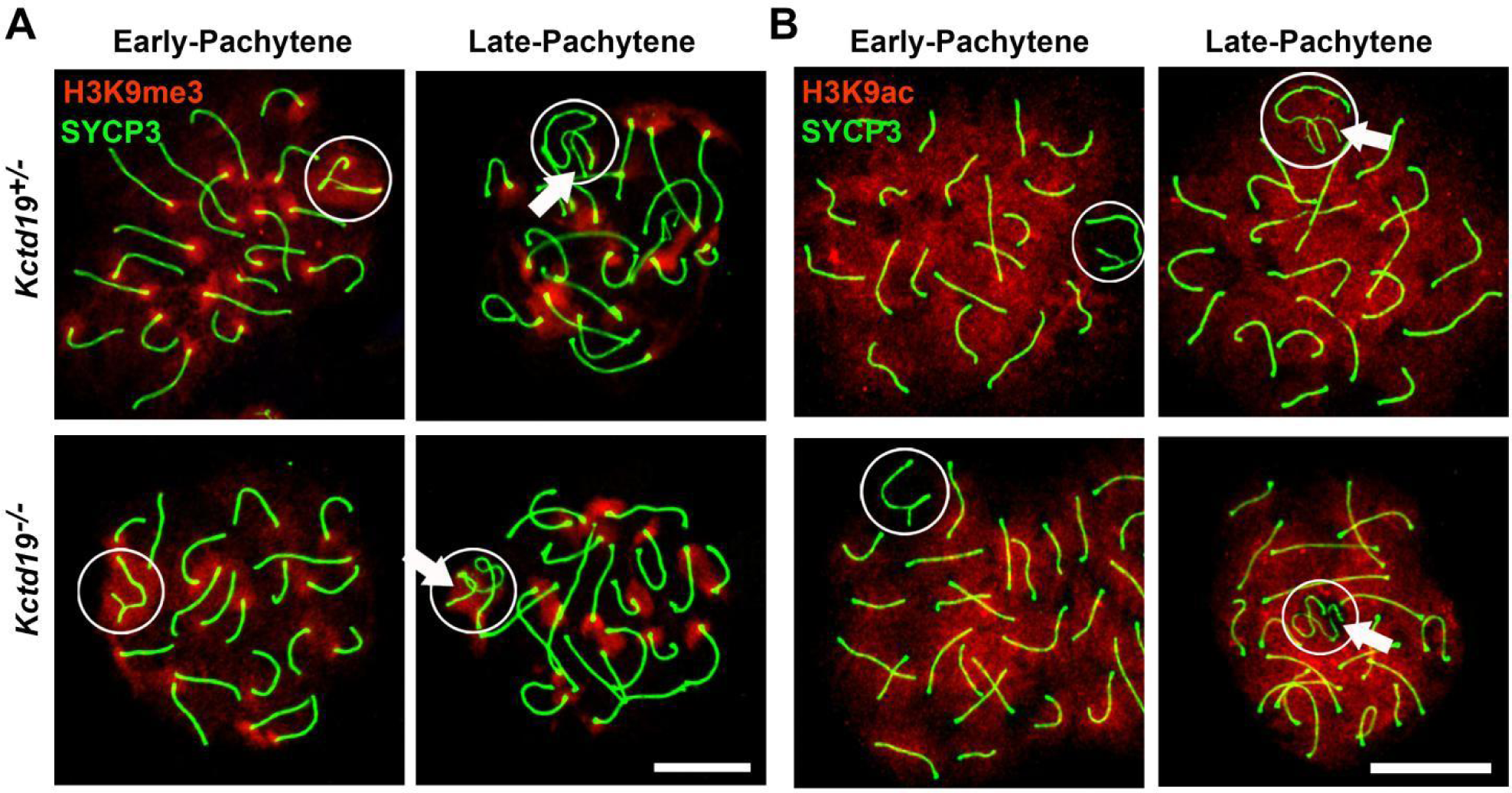
Depletion of *Kctd19* disrupts histone modifications of Y chromosome. **(A)** Immunofluorescent staining of SYCP3 and H3K9me3 on chromosome spreads of 10-week-old *Kctd19*^+/-^ and *Kctd19*^−/−^ pachytene spermatocytes. SYCP3 staining (green); H3K9me3 staining (red). Circles indicate XY bodies, and arrows indicate Y chromosomes. Scale bars: 10 μm. **(B)** Immunofluorescent staining of SYCP3 and H3K9ac on chromosome spreads of 10-week-old *Kctd19*^+/-^ and *Kctd19*^−/−^ pachytene spermatocytes. SYCP3 staining (green); H3K9ac staining (red). Circles identify XY bodies and arrows identify Y chromosomes. Scale bars: 10 μm.

**Figure 6-figure supplement 1.**
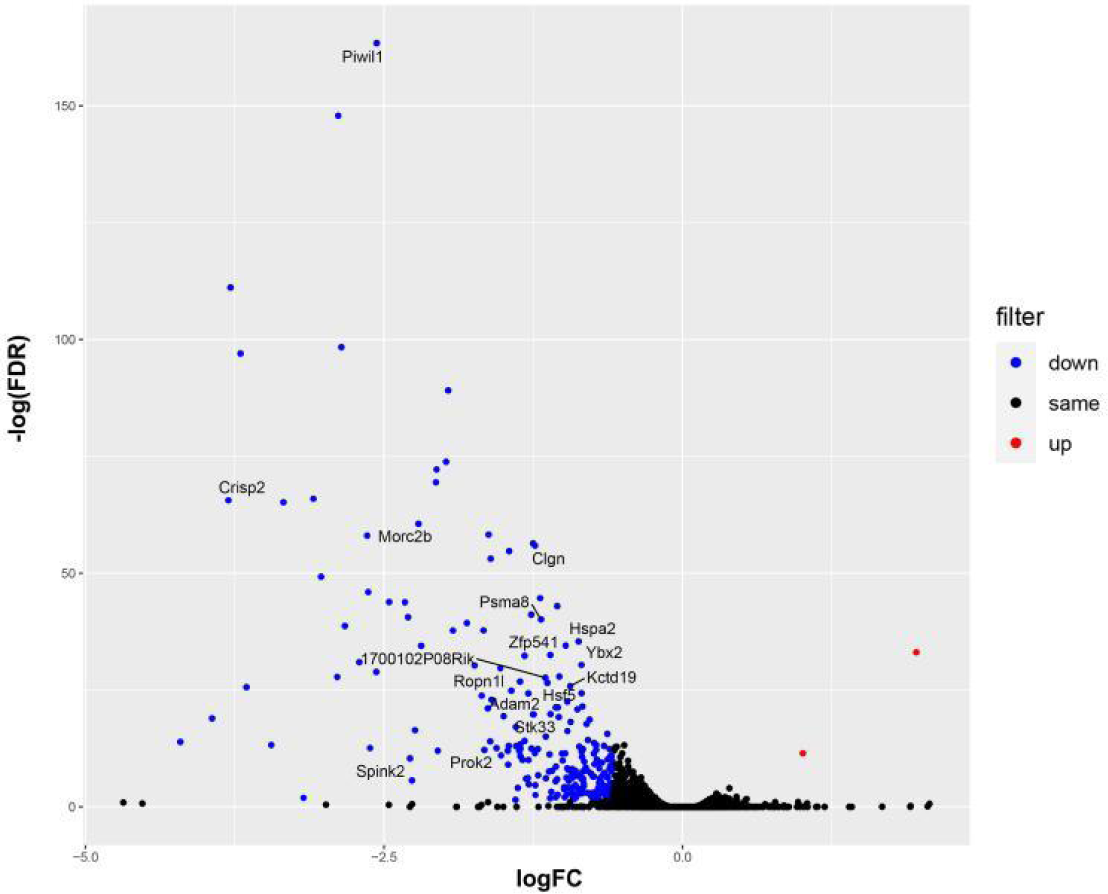
Genes exhibiting significant up- or down-regulation are showed in *Zfp541*^−/−^ testes. Volcano plot showing the level of change (log transformed normalised counts) and differential expression of genes observed between P14 *Zfp541*^+/−^ and *Zfp541*^−/−^ testes.

**Figure 6-figure supplement 2.**
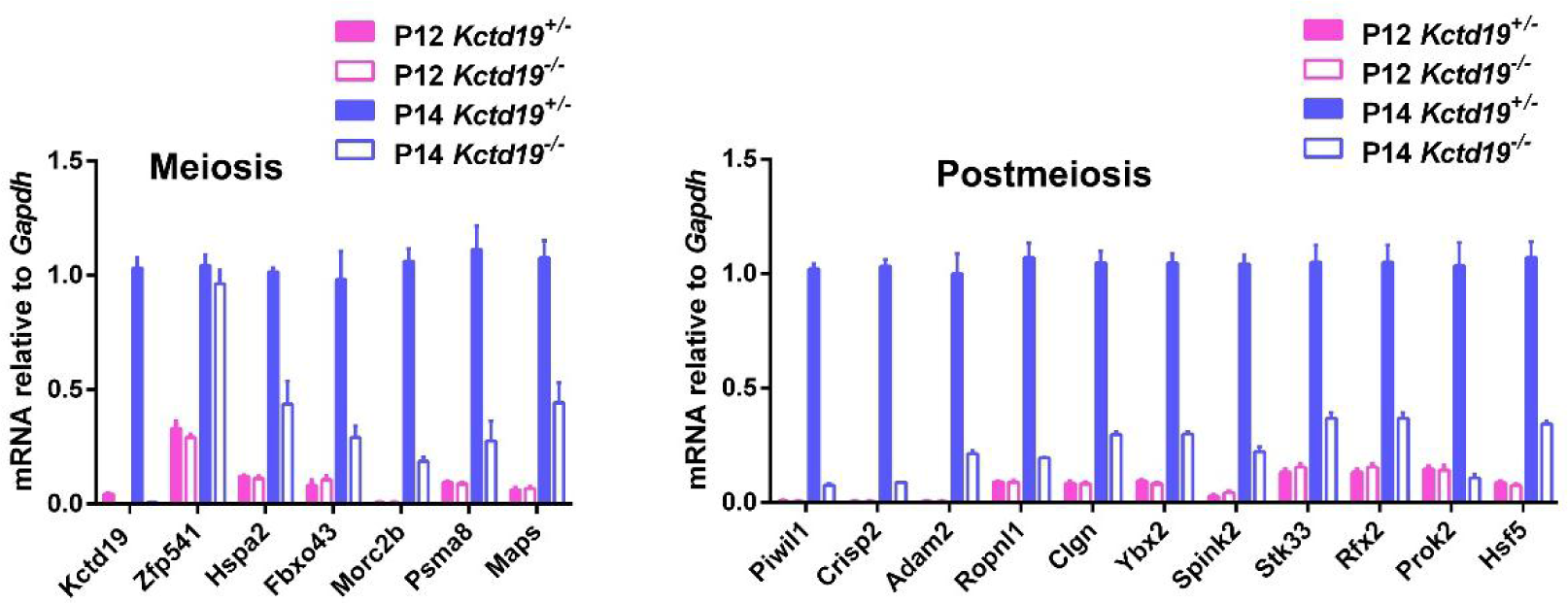
A substantial of meiotic and post-meiotic genes are downregulated in *Kctd19*^−/−^ testes, while *Zfp541 is* not affected. Validation of differentially expressed genes by qRT-PCR analysis in P12 and P14 *Kctd19*^+/-^ and *Kctd19*^−/−^ testes (n = 3, Student’s *t*-test).

